# A Genetic Screen for Temperature-sensitive Morphogenesis-defective *Caenorhabditis elegans* Mutants

**DOI:** 10.1101/2020.12.01.407221

**Authors:** Molly Christine Jud, Josh Lowry, Thalia Padilla, Erin Clifford, Yuqi Yang, Francesca Fennell, Alexander Miller, Danielle Hamill, Austin Harvey, Martha Avila-Zavala, Hong Shao, Nhah NguyenTran, Zhirong Bao, Bruce Bowerman

## Abstract

Morphogenesis involves coordinated cell migrations and cell shape changes that generate tissues and organs, and organize the body plan. Cell adhesion and the cytoskeleton are important for executing morphogenesis, but their regulation remains poorly understood. As genes required for embryonic morphogenesis may have earlier roles in development, temperature-sensitive embryonic-lethal mutations are useful tools for investigating this process. From a collection of ∼200 such *Caenorhabditis elegans* mutants, we have identified 17 that have highly penetrant embryonic morphogenesis defects after upshifts from the permissive to the restrictive temperature, just prior to the cell shape changes that mediate elongation of the ovoid embryo into a vermiform larva. Using whole genome sequencing, we identified the causal mutations in seven affected genes. These include three genes that have roles in producing the extracellular matrix, which is known to affect the morphogenesis of epithelial tissues in multicellular organisms. The *rib-1* and *rib-2* genes encode glycosyltransferases, and the *emb-9* gene encodes a collagen subunit. We also used live imaging to characterize epidermal cell shape dynamics in one mutant, *or1219*ts, and observed cell elongation defects during dorsal intercalation and ventral enclosure that may be responsible for the body elongation defects. These results indicate that our screen has identified factors that influence morphogenesis and provides a platform for advancing our understanding of this fundamental biological process.

**SUMMARY:** We performed a systematic, forward genetics screen for temperature-sensitive embryonic-lethal (TS-EL) *Caenorhabditis elegans* mutants that are specifically defective in embryonic morphogenesis. By taking advantage of temperature-upshifts, we identified several essential genes influencing morphogenesis. We also demonstrated that one mutant has defects in epidermal cell shape changes that likely account for the failure in morphogenesis. The TS-EL mutants we identified will be useful tools for advancing our understanding of the gene networks controlling cell shape changes and movements during morphogenesis.

## INTRODUCTION

Morphogenesis is a conserved process of cell shape changes and movements that ultimately organize and shape an organism. During animal embryogenesis, cell proliferation and cell fate specification produce differentiated cells, while morphogenetic changes in cell shape and position spatially organize them (Lecuit and Le Goff, 2007; Lecuit and Lenne, 2007; Sutherland, 2016). Defects in embryonic morphogenesis can result in phenotypes ranging from inconvenient to lethal. For example, morphogenetic defects are associated with vascular and neural tube closure abnormalities in humans (Greene and Copp, 2014; Ray and Niswander, 2012), and morphogenetic mechanisms are important in wound healing and cancer metastasis (Friedl and Gilmour, 2009; Martin and Parkhurst, 2004). While the cytoskeleton and cell adhesion are instrumental to cell shape changes and movements during morphogenesis, their regulation remains only partially understood.

The nematode *Caenorhabditis elegans—*with its powerful genetics, optical transparency, and a simple and invariant cell lineage—provides a powerful model system for investigating morphogenesis. The body plan of *C. elegans* is relatively simple, and elongation of the initially ovoid embryo into a vermiform larva is driven largely by epidermal cell morphogenesis. This elongation comprises three epidermal cell morphogenetic events: dorsal intercalation, ventral enclosure, and seam cell elongation (Carvalho and Broday, 2020; Chin-Sang and Chisholm, 2000; Chisholm and Hardin, 2005; Hall and Altun, 2008; Priess and Hirsh, 1986; Simske and Hardin, 2001; Sulston et al., 1983).

Most of the epidermal cells are specified by about 210-240 minutes at 20°C after the first embryonic mitosis. The resulting six rows of cells, located in a patch on the posterior dorsal side of the embryo, are organized with two inner rows (dorsal epidermal cells), two middle rows (lateral epidermal cells called seam cells), and two outer rows (ventral epidermal cells) containing 20, 20, and 18 cells, respectively. The first epidermal morphogenetic process, called dorsal intercalation, begins between 290-340 minutes, when the two rows of 10 dorsal cells on either side of the dorsal midline interdigitate. These cells become wedge-shaped and extend basolateral protrusions at their pointy ends in the direction of the dorsal midline (Figure 1A) (Williams-Masson et al., 1998). The contralateral cells then extend past each other until they make contact with seam cells on the opposite side, thus creating a single row of elongated cells along the dorsal surface (Figure 1B). Dorsal intercalation lengthens the dorsal surface relative to the lateral and ventral surfaces, creating a bean-shaped embryo (Heid et al., 2001). Actin microfilaments and microtubules are required for intercalation (Williams-Masson et al., 1998). Parallel pathways involving Rac and RhoG (Walck-Shannon and Hardin, 2016; Walck-Shannon et al., 2015), the polarity pathway members including CDC-42, PAR-2, and PAR-3 (Walck-Shannon et al., 2016), and the ephrin receptor VAB-1 (Walck-Shannon et al., 2016) control the basolateral protrusive activity for tip formation of dorsal epidermal cells undergoing intercalation. Nevertheless, the mechanisms regulating dorsal intercalation require further exploration.

**Figure 1.**
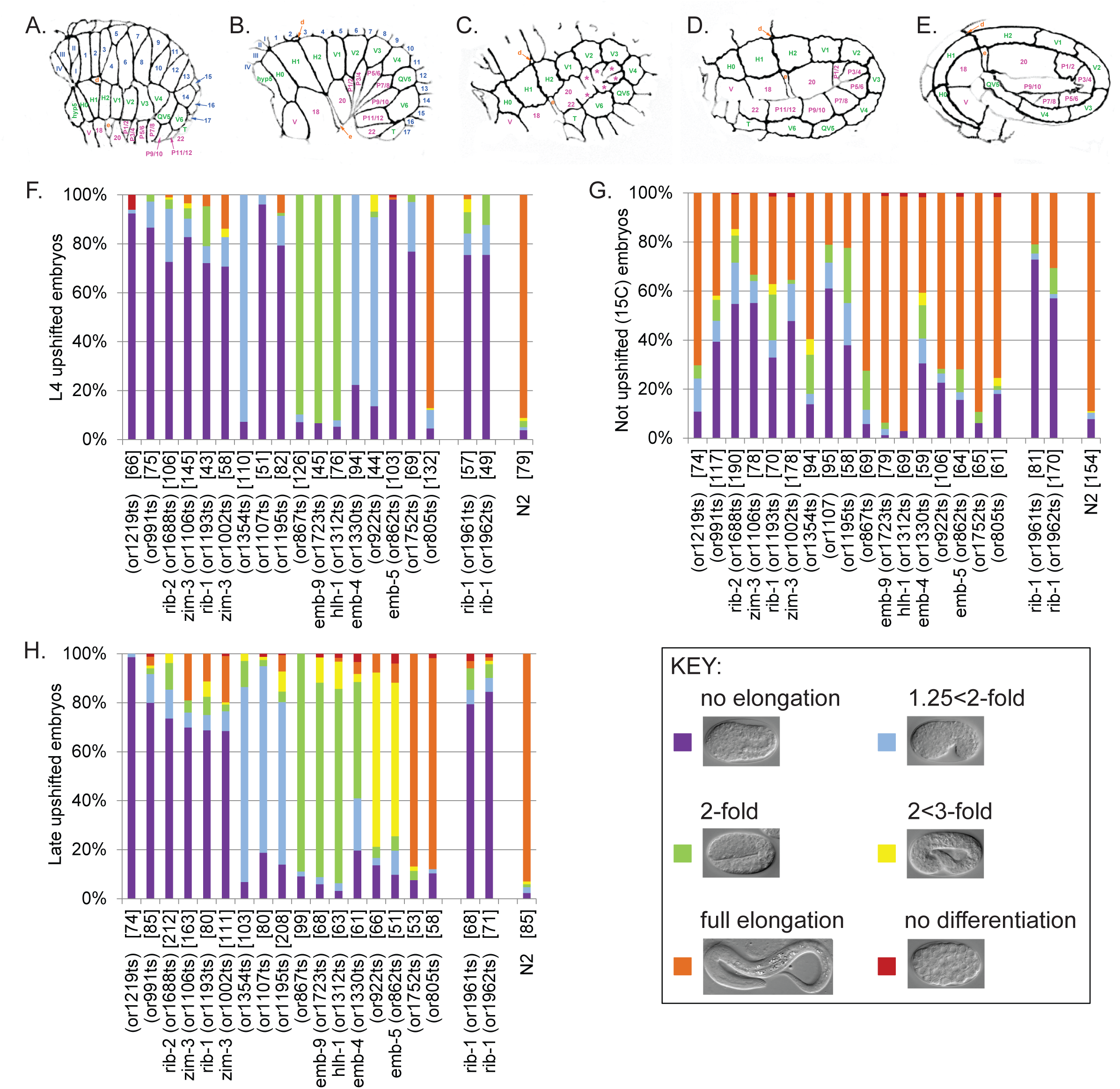
Terminal elongation-defective phenotypes for TS-EL mutants after L4 and late upshifts. **(A-E)** Schematics of wild-type embryos undergoing morphogenesis at stages just prior to bean (A), at 1.25-fold (B), at 1.5-fold (C), at 2-fold (D), and at 3- fold (E). All embryos are oriented with anterior to the left and dorsal at the top. Dorsal epidermal cell names are in blue, lateral epidermal cells (also called seam cells) in green, and ventral epidermal cells in pink. * in panel B, in order from anterior to posterior, refers to cells P1/2, P3/4, P5/6, P7/8, P9/10, and P11/12. **(F-G)** Quantification of terminal elongation-defective phenotypes for TS-EL mutant and wild-type (N2) embryos after L4 upshifts to 26°C (F), after no upshifts (G), and after late upshifts (H). Shown here are only the 17 mutants that had penetrant elongation-defective phenotypes after late upshifts (≥70% embryos exhibiting a single elongation-defective phenotype). Mutants are shown from left to right by decreasing penetrance for the most penetrant elongation category. Percent of embryos that differentiated well but arrested with no elongation (purple), or arrested at the 1.25<2-fold stage (light blue), at the 2-fold stage (green), at the 2<3-fold stage (yellow), or with full elongation (≥3-fold), or exhibited differentiation defects (red) were scored using Nomarski optics. Examples of Nomarski images of control embryos for each elongation stage are shown in the Key. The mutants are listed in the same order from left to right for both the L4 upshift (F) and no upshift (G) graphs. The CRISPR-made alleles reconstituting the *or1193*ts missense mutation in the gene *rib-1*, *or1961* and *or1962*, are included to the right of the original 17 mutants identified in the screen. If the causal mutation has been identified for a TS-EL mutant, the affected gene is listed next to the TS-EL mutant allele, and the number of embryos scored for each mutant is in brackets. See Supplemental Figure 1 for the for all 79 TS-EL mutants with penetrant elongation defects after L4 upshifts, and Supplemental Figure 2 for TS-EL mutants with low penetrance defects after late upshifts

Ventral enclosure begins as dorsal intercalation ends and serves to cover the posterior ventral surface of the embryo with a layer of epidermal cells. In the first step, two pairs of leading cells, V0 and V18 on the left side and V1 and V19 on the right side (Figure 1A), extend actin-rich filopodia towards the ventral midline until they meet and form stable adherens junctions. Regulation of actin through Rac signaling and the WAVE/SCAR and Arp2/3 complexes are required for proper ventral enclosure (Patel et al., 2008; Soto et al., 2002). Next, the ventral epidermal cells posterior to the leading cells, known as ventral pocket cells, become wedge-shaped and elongate towards the ventral midline (Figure 1B). Enclosure completes when the pocket cells meet and fuse at the ventral midline, likely by a “purse-string” mechanism utilizing an actomyosin cable similar to *Drosophila* dorsal closure (Gilbert et al., 2020; Williams-Masson et al., 1997). The movement of the ventral epidermal cells also requires the underlying neuroblasts; loss of the ephrin receptor VAB-1 in neuroblasts results in the lack of neuroblast ventral cleft closure and the subsequent non-autonomous halting of epidermal leading cell migration (Carvalho and Broday, 2020; Chin-Sang et al., 1999; George et al., 1998).

Once ventral enclosure is complete, about 350 minutes after the first cleavage, elongation of the embryo along the anterior-posterior axis begins and ends at about 600 minutes when the embryo is 4-fold in length. Elongation is divided into two stages. The second stage, which we will not discuss at length here, requires muscle twitching to elongate the embryo from 2-fold to 4-fold (Carvalho and Broday, 2020; Chisholm and Hardin, 2005). The first stage, however, is driven largely by epidermal seam cell shape changes that elongate the embryo from the bean stage to 2-fold in length. Apical actin and tubulin filament bundles are organized circumferentially in all epidermal cells and anchored at cell margins by adherens junctions (Costa et al., 1997; Costa et al., 1998; Priess and Hirsh, 1986). Active Rock/Rho only in the seam cells leads to contraction of actin and their elongation along the anterior-posterior axis (Figure 1A-E), with contractile forces generated in the seam cells conveyed to the rest of the epidermis by adherens junctions (Chan et al., 2015; Diogon et al., 2007; Gally et al., 2009; Lin et al., 2012; Martin et al., 2014; Piekny et al., 2003; Piekny et al., 2000; Wissmann et al., 1999; Wissmann et al., 1997). While regulation of the actomyosin cytoskeleton in epidermal cells, cell adhesion molecules, and intermediate-filament based tethering of the epidermis to the underlying body wall muscle cells are known to be required for these morphogenetic processes (Chisholm and Hardin, 2005), the genetic pathways that regulate them remain poorly understood.

Genetic screens in *C. elegans* have identified factors that influence morphogenesis; however, systematic and unbiased forward genetic screens designed to identify recessive, embryonic-lethal mutants with defects in morphogenesis have never been done. A dominant-negative allele of *C. elegans* Rho Kinase (ROCK; LET-502) was initially identified from a screen for zygotic lethal mutations that were defective in elongation (Wissmann et al., 1997). Additional regulators of elongation were discovered from modifier screens, starting with a suppressor screen using the dominant-negative allele of *let-502,* which led to identification of the myosin phosphatase MEL-11 (Wissmann et al., 1997) and hypomorphic and null alleles of *let-502* (Piekny et al., 2000). Similarly, the non-muscle myosin heavy chain NMY-1 was identified in a *mel-11* suppressor screen (Piekny et al., 2003). The ephrin receptor VAB-1, required for ventral enclosure, was originally identified from screens for viable mutants, with *vab-1* larva and adults displaying a notched head phenotype (Brenner, 1974; George et al., 1998). Genome-wide RNA interference (RNAi) knockdown screens have identified most of the germline-expressed genes required for early embryonic cell division and cell fate patterning (Kamath et al., 2003; Sonnichsen et al., 2005), and a limited number of genes required for embryonic morphogenesis (Diogon et al., 2007). RNAi, however, is not as effective at reducing the expression of genes required later in development, and its effectiveness varies in different cell types (Ahringer, 2006). The Auxin-inducible degron system (AID) now offers a new approach for conditional gene knockdowns (Zhang et al., 2015), but thus far this approach has not been used in large-scale unbiased screens. Moreover, AID is difficult to use during later stages of embryogenesis as the eggshell presents a barrier to the cellular uptake of auxin (Negishi et al., 2019; Zhang et al., 2015). Furthermore, while knockout consortiums have isolated deletion alleles for roughly half of the genes in *C. elegans,* many essential genes have additional requirements that complicate assessing their potential roles during morphogenesis. Temperature-sensitive embryonic-lethal (TS-EL) mutants thus provide a particularly useful tool that enables one to bypass earlier requirements and to then simultaneously reduce maternal and zygotic functions after temperature upshifts, with some genes that regulate morphogenesis having both maternal and zygotic contributions (Piekny et al., 2000).

To identify additional genes required for embryonic morphogenesis, we have examined a collection of about 200 TS-EL mutants with normal early embryonic cell divisions. 17 of these mutants appear to make roughly normal numbers of differentiated cell types but arrest with partial or no elongation after upshifts to the restrictive temperature just prior to the beginning of embryonic elongation. Using a combination of classical genetics and whole genome sequencing, we have identified the causal mutations in eight of these mutants, affecting seven genes. These include alleles of *rib-1* and *rib-2*, which encode heparan sulfate synthesis proteins, and *emb-9*, which encodes an extracellular matrix collagen protein, all genes that are known to have roles in morphogenesis in other systems (Clause and Barker, 2013; Daley and Yamada, 2013; Jayadev and Sherwood, 2017; Poulain and Yost, 2015; Rozario and DeSimone, 2010). We also examined the epidermal cell shape changes that normally occur during elongation in the mutant *or1219*ts and identified defects in dorsal intercalation and ventral enclosure. These results provide the foundation for extending our understanding of the gene networks that regulate and execute embryonic morphogenesis in *C. elegans*.

## MATERIALS AND METHODS

### *C. elegans* strains and culture

Temperature-sensitive embryonic-lethal (TS-EL) mutants in a *lin-2*(*e1309*) background were isolated after chemical mutagenesis with N-ethyl-N-nitrosourea (ENU), as previously described (Encalada et al., 2000; Kemphues et al., 1988; O’Rourke et al., 2011). The 191 TS-EL mutants screened in this current study are listed in Table S1. The mutants of known genes used for complementation tests are listed in Table S2. The *C. elegans* Bristol strain (N2) was used as the wild-type strain, the Hawaiian strain (CB4856) was used for SNP mapping, and FT63 *xnIs17[dlg-1p::DLG-1::GFP, rol-6*(*su1006*)] X was used to mark epidermal cell membranes (CGC). All strains were maintained at the permissive temperature (15°C) on standard nematode growth medium plates seeded with the *E. coli* strain OP50 (Brenner, 1974).

### CRISPR

The appropriate sgRNA and PAM sites were selected using the website http://crispr.mit.edu/ to recreate the *rib-1(or1193*ts) point mutation: cAt to cTt equating to the amino acid change H126L. The injection mixture was prepared according to Dokshin et al (2018) and injected into young N2 adults: *rib-1* repair oligo ssDNA (5’ATTGGATCCGTCAGTTTGGAATAATGGAAGAAATC**<u>T</u>**TCTGAT*C*TT*T*AA*C*TT*T*TA*<u>C</u>*<u>C A*T*GG</u>AACTTTTCCTGATTATGATGATCATAATTTAAA; ***<u>or1193</u>*<u>ts point mutation</u>**, *silent mutations*, <u>NcoI restriction enzyme site</u>; IDT), crRNA (5’CTGATTTTCAATTTCTATCA; IDT), trRNA (IDT), Cas9 RNP (IDT), and the co-injection marker *rol-6(su1006)* on plasmid PRF4. The F_1_ progeny of the injected animals were selected for the roller phenotype and screened by PCR (forward primer 5’ TTGGAAGTGTTCGGTTACAAAA; reverse primer 5’ AAACTAAAATTGGCACGAAACG; IDT) and NcoI restriction digestion (New England Biolabs). Non-roller, homozygous mutant worms were identified and outcrossed to N2 and the point mutation was confirmed by Sanger sequencing (Sequetech). Two independent lines, EU3219 *rib-1*(*or1961*ts) and EU3220 *rib-1*(*or1962*ts), were analyzed for embryonic lethality and terminal phenotypes after upshift.

### Temperature-sensitive embryonic-lethality assay

Embryonic lethality was scored by singling out ten L4 hermaphrodites onto individual plates and incubating them at the permissive temperature (15°C) for two nights and the restrictive temperature (26°C) for one night. Parents were then removed and the number of eggs and larvae were counted. Plates were returned to the same temperature for two more nights at 15°C or one more night at 26°C and unhatched eggs were counted to determine the percent non-hatching rate.

### Terminal phenotyping assay

#### L4 Upshifts

Homozygous mutant worms were raised at the permissive temperature (15°C) on agar plates seeded with bacteria until the last larval stage (L4) and then up-shifted to the restrictive temperature (26°C) for a minimum of eight hours. Early stage embryos (1-8 cell stages) were then dissected and sorted from young gravid adults in a droplet of M9 solution and transferred with a mouth pipette to a 4% agar pad on a glass slide, overlaid with a coverslip and sealed with 100% petroleum jelly to prevent desiccation. Embryos were then allowed to develop for an additional 15-16 hours at 26°C, at which time they were examined with a compound Zeiss microscope using Nomarski optics. Embryos were scored for arrest at different elongation stages: no elongation, 1.25<2-fold, 2-fold, 2<3-fold, and full elongation (>3-fold). Embryos were also assessed for differentiation by having roughly normal morphology of cells, gut granules, neuronal processes, twitching, or lack of obviously undifferentiated cells using DIC and polarizing filters. A penetrant phenotype was defined as ≥70% of embryos displaying a single elongation stage, and variable phenotypes were defined as 40-70% of embryos displaying a single elongation stage.

#### Late Upshifts

Homozygous mutant worms were raised at the permissive temperature (15°C) on agar plates seeded with bacteria until they became young gravid adults. Early stage embryos (1-8 cell stages) were dissected and sorted at 15°C. Slides were prepared as described above for “L4 upshifts”, but slides were maintained at 15°C for 7-7.5 hours before being upshifted to the restrictive temperature (26°C) for 15-16 hours. Embryos were then examined and scored using Nomarski optics as described above for “L4 upshifts”. The 7-7.5 hours (420-450 minutes) at 15°C corresponds to ∼five hours at 20°C (∼300 minutes) at which point embryos have bypassed most of the cell division and cell fate patterning developmental programs and are at a stage just prior to the morphogenetic programs of dorsal intercalation, ventral enclosure, and elongation (Chisholm and Hardin, 2005; Sulston et al., 1983).

### Genetic Analysis

TS-EL mutants with penetrant terminal phenotypes after L4 upshift were outcrossed to N2 males to remove the *lin-2*(*e1309*) mutation present in the parental strains as previously described (Lowry et al., 2015). Concurrently, we assessed if the mutations appear recessive. We expected 0-25% embryonic lethality in the broods of 10 individualized TS-EL/+ F_1_ heterozygotes after shifting to the restrictive temperature, with 0% expected for recessive maternal-effect mutants and 25% for recessive zygotic-effect mutants. Second, we determined if the mutations demonstrate Mendelian segregation. When lethality is due to a single gene mutation, 25% of the 80 individualized F_2_ worms produced after F_1_ TS-EL/+ self-fertilization are expected to make broods of dead embryos at the restrictive temperature, whereas if lethality is due to a double gene mutation, 6.25% of F_2_ worms are expected to make broods of dead embryos. TS-EL mutants that appeared dominant (>∼25% embryonic lethality) or a multiple gene mutation (≤10% F_2_’s with embryonic lethal broods) were removed from further analysis.

### Identification of causal mutations

We performed whole genome sequencing, SNP mapping, and sequencing analysis as described previously (Lowry et al., 2015). Briefly, homozygous TS-EL mutants were mated to a polymorphic Hawaiian strain, CB4856, with SNPs roughly every ∼1.0 kb compared to the N2 parental strain. 10-30 homozygous TS-EL F_2_ strains were then isolated from each cross and genomic DNA was pooled for sequencing using the Qiagen DNAeasy and Kapa HyperPlus Preparation Kits. Whole genome sequencing was performed using paired-end NextSeq 500 Illumina sequencing in the Genomics and Cell Characterization Core Facility (GC3F) at the University of Oregon. The data files were then processed using the Galaxy platform as previously described (Lowry et al., 2015). TS-EL mutations were mapped to intervals of a few megabases and candidate mutations were identified within those intervals, filtering for missense and splice-site mutations in coding sequences of genes known to be essential based on genome-wide RNAi screens or previous mutagenesis screens. Candidate genes were then tested using complementation tests with known alleles when available (Table S2). Homozygous mutations were visualized using the Integrative Genomics Viewer from the Broad Institute (Robinson et al., 2017; Robinson et al., 2011; Thorvaldsdottir et al., 2013).

### Assessing cell shape changes in live embryos

To investigate the cell shape changes and movements that mediate early elongation, *xnIs17[dlg-1p::DLG-1::GFP, rol-6*(*su1006*)] X that marks epidermal cell membranes (Firestein and Rongo, 2001) was introduced by mating into the *or1219*ts mutant background. Two strains were produced and used for analysis: (1) EU3187 *or1219*ts I; *ruIs32*[*unc-119*(*+*) *pie-1p::GFP::H2B*] III; *xnIs17[dlg-1p::DLG-1::GFP, rol-6*(*su1006*)] X, and (2) EU3208 *or1219*ts I; *xnIs17[dlg-1p::DLG-1::GFP, rol-6*(*su1006*)] X. As both strains yielded the same results, data collected from both were combined for our analysis of phenotypes. To assess epidermal cell shapes in live embryos, 2-cell stage control (*xnIs17*) or mutant (*or1219*ts *xnIs17*) embryos were harvested and sorted from gravid adults at the permissive temperature (15°C). The embryos were allowed to develop in a watch glass in M9 for six hours at 15°C, at which point, the embryos were mounted on a 2% agarose pad sandwiched between a 22×40mm coverslip and an 18×18mm coverslip sealed with 100% petroleum jelly to prevent desiccation. We used the Cherry Temp system from Cherry Biotech, a fluidics device that permits temperature control of biological samples while imaging at microscopes. The mounted embryo coverslip sandwich was attached with tape to the Cherry Temp fluidics chip set to 15°C. At 6.5 hours after harvest, the late stage embryos were either left at the control temperature of 15°C for two hours or upshifted to 26°C for one hour to age them to bean stage, when DLG-1::GFP is expressed in the embryo (Firestein and Rongo, 2001).

After the embryos were aged to bean stage (time 0 minutes), embryos were then imaged every 40 minutes at 15°C or every 20 minutes at 26°C for several hours using an Andor Leica spinning-disc confocal microscope. Since embryonic development is twice as fast at 26°C compared to 15°C (Corsi et al., 2015), the aging and imaging times were adjusted to allow for direct comparisons of epidermal cell shapes between the two temperatures for each imaging time point. Entire embryos were imaged by at 100x by acquiring multiple stacks (≥50 slices) with a step size of 0.5 μm, 488 laser power of 0.5%, EM Gain of 300, offset of 0, and an exposure time of 100 ms. Maximum projection images were created using ImageJ and converted from 16 bit to 8 bit. Contrast (output levels, gamma) were adjusted to optimize the visibility of the epidermal cell membranes and figures were built in Adobe Photoshop CS5 and Adobe Illustrator CS5.

### Data and Reagents

Strains will be made available to the Caenorhabditis Genetics Center (CGC) upon publication (https://cgc.umn.edu/). The authors affirm that all data necessary for confirming the conclusions of the article are present within the article, figures, and tables.

## RESULTS

### Identifying recessive, single locus, temperature-sensitive embryonic-lethal mutations that confer penetrant elongation defects

To identify essential genes required for embryonic morphogenesis, we examined a collection of previously isolated temperature-sensitive embryonic-lethal mutants (Encalada et al., 2000; Lowry et al., 2015; O’Rourke et al., 2011), focusing on 191 that appeared to have normal early embryonic cell divisions but nevertheless failed to hatch. We initially examined the terminally differentiated phenotypes of embryos produced after shifting L4 larvae to the restrictive temperature (see Materials and Methods), to identify mutants with roughly normal patterns of differentiated cell types that failed to elongate. 79 mutants produced embryos with penetrant defects (>70% arresting with roughly the same extent of embryonic elongation stage), and 30 mutants were more variable (40-70% arresting with roughly the same extent of embryonic elongation) (Figure S1). Of the 79 mutants with penetrant defects, the majority arrested with no elongation (63); the remainder (16) arrested with some elongation but failed to hatch (Figures 1F, S1A-B).

The large number of mutants with penetrant elongation defects suggested to us that many of these mutants might have earlier defects in cell fate specification or cell division that indirectly affect morphogenesis. We therefore next raised mutant worms to adulthood and allowed their embryos to develop at the permissive temperature (15°C) to bypass the bulk of proliferation, cell fate specification and differentiation, before upshifting embryos to the restrictive temperature (26°C) shortly before the morphogenetic elongation processes of dorsal intercalation, ventral enclosure, and elongation (see Materials and Methods). After these “late upshifts,” the embryos produced by most of the mutants elongated and hatched (Figure S2), indicating that either earlier requirements during embryogenesis were responsible for the elongation defects we observed after L4 larval upshifts, or that the TS mutations were insufficiently fast-acting to impair morphogenesis. However, embryos from 17 mutants still failed to elongate after these late upshifts (Figure 1H), suggesting that the affected loci are more likely to have direct roles in morphogenesis. All of these 17 mutants displayed a greater extent of elongation and hatching when kept at 15°C continuously (Figure 1G). Of the 63 mutants that were penetrant for no elongation after L4 upshifts, only six remained penetrant after late upshifts. Of the 16 mutants that were penetrant for partial elongation after L4 upshifts, 11 mutants remained penetrant for partial elongation after later upshifts. We conclude that these 17 mutants are more likely to be directly involved in the regulation or execution of embryonic morphogenesis.

We next genetically characterized the 79 mutants with penetrant elongation defects to determine if the mutations appeared recessive and affected a single locus (see Materials and Methods). All 17 mutants with penetrant elongation defects after late upshifts appeared to be recessive and affect a single locus. All but four of the remaining 62 mutants with penetrant elongation defects only after L4 upshifts appeared to be recessive, and all but three appeared to affect a single locus (Tables 1, S3). The seven mutants that were either semi-dominant or appeared to affect more than a single locus were not further analyzed.

**Table 1.** Embryonic lethality and genetic characterization of penetrant late upshifted TS-EL mutants.

| TS allele | % EL at 15°C (n) | % EL at 26°C (n) | Heterozygous<br>% EL at 26°C (n) | Segregation<br>frequency (n) |
| --- | --- | --- | --- | --- |
| *or805* | 1.3 (388) | 99.4 (882) | 19.2 (1140) | 28.8 (80) |
| *or862* * | 0.6 (334) | 98.4 (382) | 1.1 (263) | 22.5 (80) |
| *or867* | 3.9 (332) | 100.0 (565)<sup>‡</sup> | 2.7 (594) | 22.5 (80) |
| *or922* | 5.0 (317) | 98.9 (725) | 21.3 (2004) | 28.9 (90) |
| *or991* | 28.9 (516) | 77.0 (575) | 0.8 (384) | 20.0 (80) |
| *or1002* * | 66.9 (662) | 91.7 (945) | 8.5 (863) | 27.8 (90) |
| *or1106* * | 63.7 (870) | 95.8 (402) | 2.1 (368) | 16.5 (79) |
| *or1107* | 0.2 (543) | 97.2 (1008) | 27.1 (1357) | 32.9 (82) |
| *or1193* * | 28.2 (1613) | 95.1 (2607) | 13.9 (1365) | 21.8 (78) |
| *or1195* | 74.4 (273) | 97.6 (252) | 6.7 (312) | 34.6 (78) |
| *or1219* | 12.7 (900) | 99.7 (651) | 8.3 (229) | 18.8 (80) |
| *or1312* * | 1.7 (631) | 99.4 (1167) | 16.6 (465) | 17.7 (79) |
| *or1330* * | 50.0 (1205) | 100.0 (916) | 27.2 (687) | 23.0 (100) |
| *or1354* | 69.0 (507) | 100.0 (495) | 2.8 (435) | 23.6 (89) |
| *or1688* * | 56.6 (806) | 87.7 (929) | 2.7 (1134) | 19.3 (88) |
| *or1723* * | 1.5 (390) | 100.0 (423) | 16.8 (285) | 21.3 (80) |
| *or1752* | 0.9 (318) | 100.0 (548) | 5.8 (452) | 26.3 (80) |
| *or1961* | 22.9 (772) | 95.4 (1621) | 18.0 (1357) | 22.4 (49) |
| *or1962* | 23.3 (848) | 95.9 (1959) | 19.4 (1150) | 20.0 (50) |

\* = causal mutation identified
<sup>‡</sup> = embryonic lethality and PAT phenotypes combined (22.3 and 77.7, respectively)

### Identifying causal mutations

After identifying likely morphogenesis-defective mutants, we next sought to identify the causal mutations in both the 17 penetrant late stage upshifted mutants and some of the 62 mutants that were penetrant only after L4 upshifts. To identify candidate mutations, we used a whole genome sequencing approach combined with genome-wide Single Nucleotide Polymorphism (SNP) mapping (Doitsidou et al., 2010; Lowry et al., 2015; Minevich et al., 2012). Thus far, we have completed whole genome sequencing for 37 of the 62 mutants that were penetrant for elongation defects only after L4 upshifts, and for 14 of the 17 mutants that had penetrant elongation defects after late upshifts (data not shown; Figures 2A, S3A, S4A-B, S5A, S6A, S7A, S8A). After identifying the region of interest, typically spanning several megabases, we focused further effort on missense and splice site mutations in essential genes, as we have found that these types of mutations account for most of the TS alleles we have isolated (Lowry et al., 2015; O’Rourke et al., 2011).

**Figure 2.**
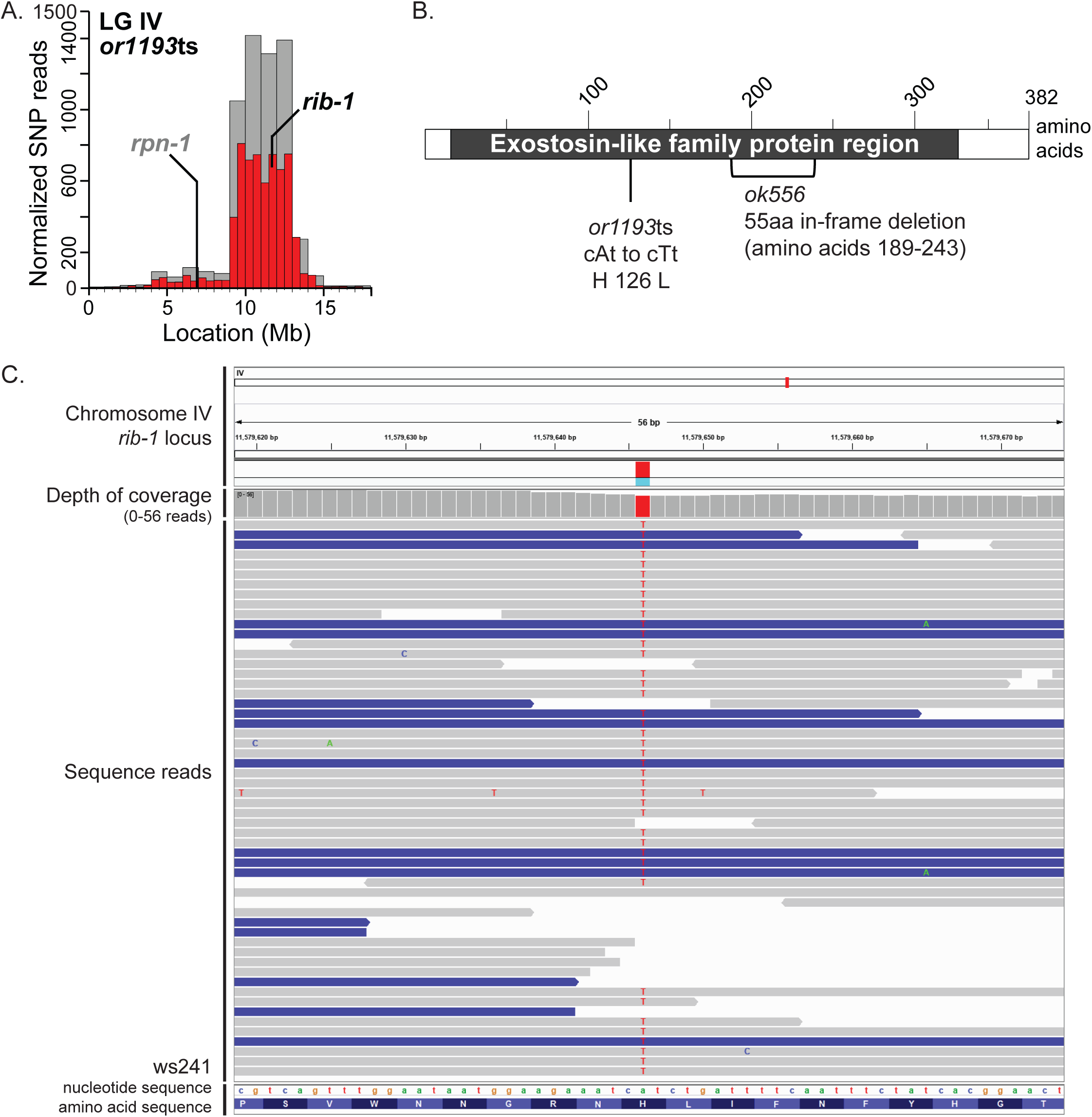
Identification of *rib-1(or1193*ts) causal mutation. **(A)** SNP mapping data for the *or1193*ts mutant on linkage group IV with identified causal mutations, showing the frequency of homozygous parental alleles plotted against chromosomal position in bins of either 1 megabase (gray bars) or 0.5 megabase (red bars). Candidate essential genes in which missense mutations were detected are indicated. For complementation test in which *or1193*ts failed to complement the known genetic mutation, the gene is dark and bolded; for complementation test in which *or1193*ts complemented the known mutation, the gene is gray and bolded. See Table 2 for complementation test results. **(B)** Sequence alterations in the predicted RIB-1 protein, isoform a, for *or1193*ts and *ok566,* the known mutant allele used for the complementation test. RIB-1 is orthologous to human exostosin-like glycosyltransferase 2 (EXT2) (Kitagawa et al., 2007) and shares sequence homology to the Exostosin family (black box corresponding to amino acids 17-333; hmmpanther, INTERPRO, and Pfam on https://wormbase.org). **(C)** Integrative Genomics Viewer (Broad Institute) screenshot of the sequencing reads at the site of the *or1193*ts missense mutation at the *rib-1* locus. The red bar in the depth of coverage section indicates the homozygosity of the A to T nucleotide change across the reads. Gray lines indicate all bases matched the reference sequence; blue lines imply reads of the opposite strand, (https://software.broadinstitute.org/software/igv/interpreting_pair_orientations). Single nucleotide changes are indicated on each read (green A, blue C, orange G, and red T). Nucleotide and amino acid sequences are read from left to right.

**Table 2.** Causal mutation identification for penetrant late upshifted TS-EL

| TS allele | gene | Tested allele | Transheterozygote<br>% EL at 26°C (n) |  |
| --- | --- | --- | --- | --- |
| *or862* | ***emb-5*** | *hc61ts* | 100.0 | (266) |
|  | *mua-3* | *rh195ts* | 4.2 | (1311) |
| *or1002* | ***zim-3*** | *or1106ts* | 95.8 | (2211) |
| *or1106* | ***zim-3*** | *tm2303ts* | 97.1 | (208) |
|  | *rpn-1* | *ok2259* | 11.8 | (561) |
| *or1193* | ***rib-1*** | *ok556* | 97.1 | (1518) |
|  | *rpn-1* | *ok2259* | 2.7 | (990) |
| *or1312* | ***hlh-1*** | *cc561ts* | 100.0 | (504) |
| *or1330* | ***emb-4*** | *hc60ts* | 100.0 | (309) |
| *or1688* | ***rib-2*** | *gk318* | 97.8 | (1221) |
|  | *dcr-1* | *ok247* | 16.4 | (860) |
| *or1723* | ***emb-9*** | *hc70ts* | 99.9 | (1188) |
|  | *zfp-1* | *ok544* | 51.8 | (326) |
| *or1961* | ***rib-1*** | *or1193ts* | 90.4 | (1470) |
| *or1962* | ***rib-1*** | *or1193ts* | 93.4 | (1520) |

**Bold** denotes the known mutation failed to complement the TS-EL allele

We next performed complementation tests with known mutant alleles to identify causal mutations (see Materials and Methods). Of the 17 mutants with penetrant elongation defects after late upshifts, eight failed to complement previously isolated mutations in essential genes while complementing mutations in other candidate genes; the causal mutations have yet to be identified for the other nine mutants (Table 2). We confirmed the homozygosity of each causal mutation by viewing the sequencing reads at the site of the mutation using the Integrative Genomics Viewer from the Broad Institute (Figures 2C, S3C, S4D, S5C, S6C, S7C, S8C) (Robinson et al., 2017; Robinson et al., 2011; Thorvaldsdottir et al., 2013). The eight confirmed causal mutations represent seven different genes (Tables 2, S4).

Three of the causal mutations we have identified map to genes that are likely to directly influence morphogenesis. The alleles *or1193*ts and *or1688*ts failed to complement previously isolated mutations in *rib-1* and *rib-2*, respectively, which function in the same protein modification pathway (Table 3). They encode heparan sulfate synthesis proteins that are known to regulate morphogenesis in *C. elegans* and other organisms (Franks et al., 2006; Kitagawa et al., 2001; Kitagawa et al., 2007; Morio et al., 2003; Poulain and Yost, 2015), although how post-translational sugar modifications influence morphogenesis is still unclear (see Discussion). We also identified *or1723*ts as a mutation in *emb-9*, which encodes a collagen subunit (Table 3), a structural component of extracellular matrices which are known to provide support, both molecularly and mechanically, to cells undergoing morphogenesis (Clause and Barker, 2013; Daley and Yamada, 2013; Guo and Kramer, 1989; Jayadev and Sherwood, 2017; Rozario and DeSimone, 2010). Therefore, our screen appears to have identified mutations in genes that directly influence morphogenesis.

**Table 3.** Gene orthologs and functions for late upshifted causal TS-EL mutations.

| Gene | TS allele | Human ortholog | Function |
| --- | --- | --- | --- |
| **Cell Division** (meiosis and/or mitosis) |  |  |  |
| *zim-3* | *or1002*<br>*or1106* | none | Zinc-finger in meiosis |
| **Gene Expression** (signaling/transcription factors) |  |  |  |
| *emb-5* | *or862* | SPT6 | RNA polymerase II transcription elongation factor |
| *hlh-1* | *or1312* | MRF | Basic helix-loop-helix transcription factor |
| **Gene Expression** (nucleosome/chromatin regulation) |  |  |  |
| *emb-4* | *or1330* | Aquarius | Nuclear protein with putative AAA ATPase domain |
| **Extracellular Matrix** |  |  |  |
| *emb-9* | *or1723* | COL4A3, 5, 6 | Collagen |
| **Protein Modification** |  |  |  |
| *rib-1* | *or1193* | EXT1 | GlcNAc transferase for synthesis of heparan sulfate proteoglycans |
| *rib-2* | *or1688* | EXTL3 | alpha 1,4-N-acetylglucosaminyltransferase |

We also performed complementation tests with several mutants that exhibited penetrant elongation defects only after L4 upshifts. 24 of these mutant alleles failed to complement previously isolated mutations in 14 genes (Table S5). All the mutations were missense in nature except for one allele: *or1237*ts is a nonsense mutation in gene *fntb-1* (Table S6). Most of these genes encode proteins with general roles in gene expression (Table S7), consistent with our hypothesis that L4 upshifts might lead to low penetrance or late defects in cell division or cell fate specification that indirectly interfere with elongation. For example, we identified alleles of *glp-1,* which encodes a Notch receptor involved in blastomere specification (Good et al., 2004); *chaf-1*, which encodes a chromatin assembly factor subunit (Nakano et al., 2011; Shibata et al., 2019); and *zwl-1*, which encodes a kinetochore protein (Gassmann et al., 2008).

While complementation tests are useful for identifying causal mutations, they are not fully conclusive. For example, other mutations in the two genetic backgrounds might interact to generate the observed failure to complement. Furthermore, while six of the eight causal mutations we identified in strains that exhibited penetrant elongation defects after late upshifts are missense mutations, the *or1106*ts strain has a nonsense mutation at codon 199 in the gene *zim-3*, and the *or1330*ts strain has 1 bp insertion at codon 967 that results in a frameshift and early stop at codon 1004 in the gene *emb-4* (Table S4; Figures S4C, S6B). Such mutations are less likely than missense mutations to result in conditional embryonic lethality. To more rigorously determine if a complementation test correctly identified the causal mutation, we used CRISPR genome editing to introduce the *or1193*ts missense mutation into the *rib-1* gene, using the N2 wild-type strain background. We independently isolated two alleles, *or1961* and *or1962*, that both introduce the same amino acid change as *or1193*ts, an A to T missense mutation that substitutes a histidine for a leucine at codon 126 in the *rib-1* gene (Figure 2B; Table S4; see Materials and Methods). Both *or1961* and *or1962* behaved as recessive single locus mutations and exhibited conditional embryonic lethality (Table 1), and both resulted in elongation defects that resembled those observed with the *or1193*ts mutant, after both L4 larval and later embryonic temperature upshifts (Figure 1F-H). Furthermore, both CRISPR alleles failed to complement the original *or1193*ts mutant (Table 2). These results support our conclusion that the *or1193ts* missense mutation in *rib-1* is responsible for the observed conditional embryonic lethality and morphogenesis defect.

### Epidermal cell shape changes that mediate embryonic elongation are defective in *or1219*ts mutant embryos

To further explore whether the 17 TS-EL mutants with penetrant elongation defects after late upshifts are specifically defective in embryonic morphogenesis, we used live imaging to analyze epidermal cell shape changes in *or1219*ts, which exhibited the most highly penetrant elongation defect. To assess epidermal cell shapes over time (see Materials and Methods), we used genetic crosses to introduce *or1219*ts into a transgenic strain expressing a GFP fusion to DLG-1, which marks epidermal cell membranes (Firestein and Rongo, 2001). We first examined *or1219*ts mutant embryos kept at 15°C and observed that their development was delayed relative to control embryos. By 400 minutes after the start of imaging, nearly all control embryos had elongated to ≥3-fold in length, while about 60% of *or1219*ts embryos had elongated ≤2- fold in length (Figures 3, 5A). 70% *or1219*ts embryos did elongate to ≥3-fold by 720 minutes and were still alive after the extended period of imaging (Figures 3, 5A-B). We also observed that after the late temperature upshifts prior to elongation, epidermal cell fate patterning was relatively normal in *or1219*ts embryos. The number, shape, and location of epidermal cells in about 90% of upshifted *or1219*ts mutant embryos appeared similar to control embryos (Figures 4, 5A,). As a few *or1219*ts embryos displayed abnormal organization of the epidermal cells even when kept at 15°C (∼6%; Figure 5A), there may be a low level of cell fate patterning defects in this mutant background.

**Figure 3.**
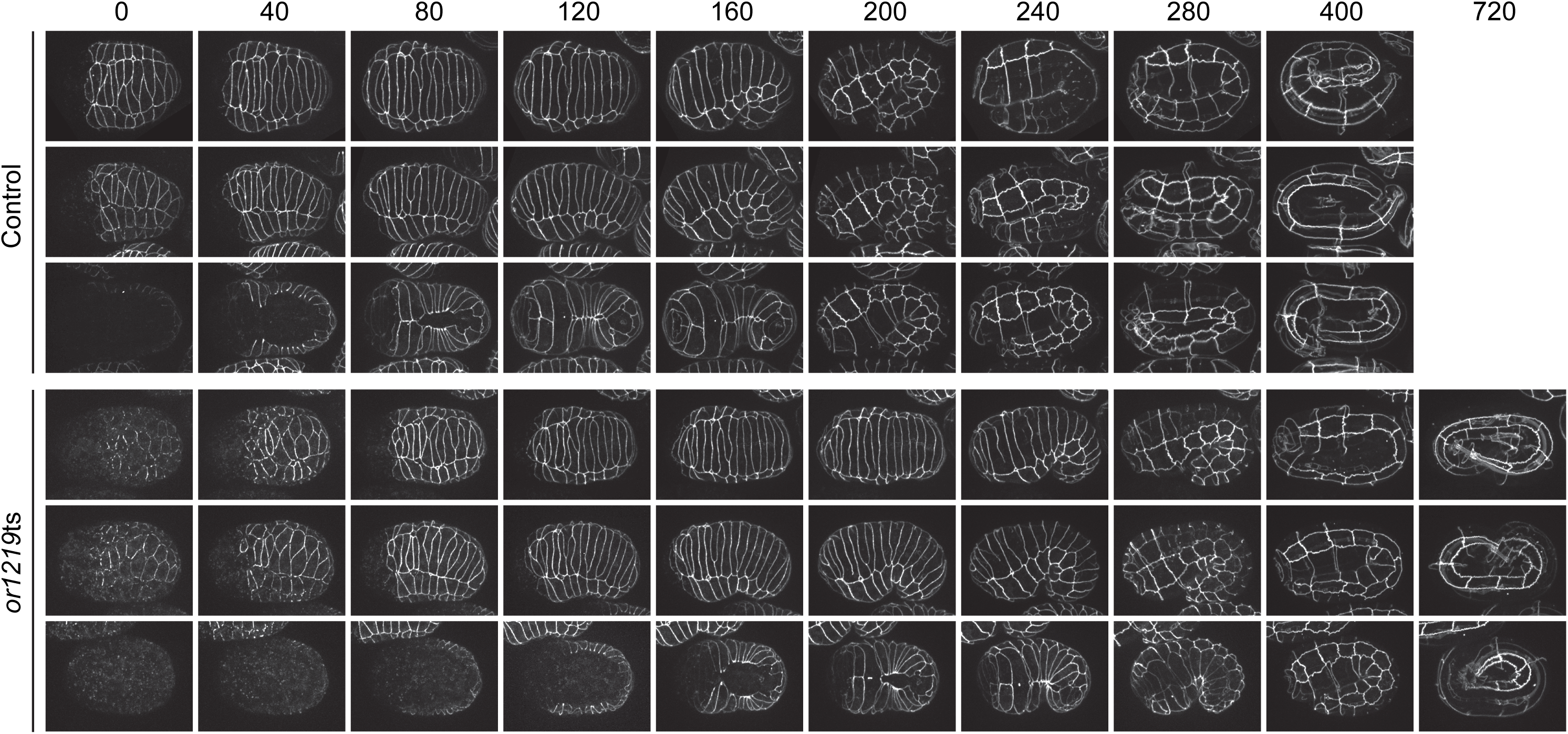
*or1219*ts embryos have delayed development at the permissive temperature but still elongate. Maximum projection images of live control (top 3 rows) or mutant (*or1219*ts; bottom three rows) embryos, expressing DLG-1::GFP to mark epidermal cell membranes, kept at 15°C. Minutes are listed across the top; time 0 corresponds to the start of imaging at the bean stage (two hours at 15°C after the late upshift time point; see Materials and Methods). Embryos were imaged every 40 minutes for 400 minutes (control embryos) or 720 minutes (*or1219*ts mutant embryos). Each row represents a single representative embryo over time in the dorsal orientation (top rows; 1 and 4), lateral orientation (middle rows; 2 and 5), and ventral orientation (bottom rows; 3 and 6).

**Figure 4.**
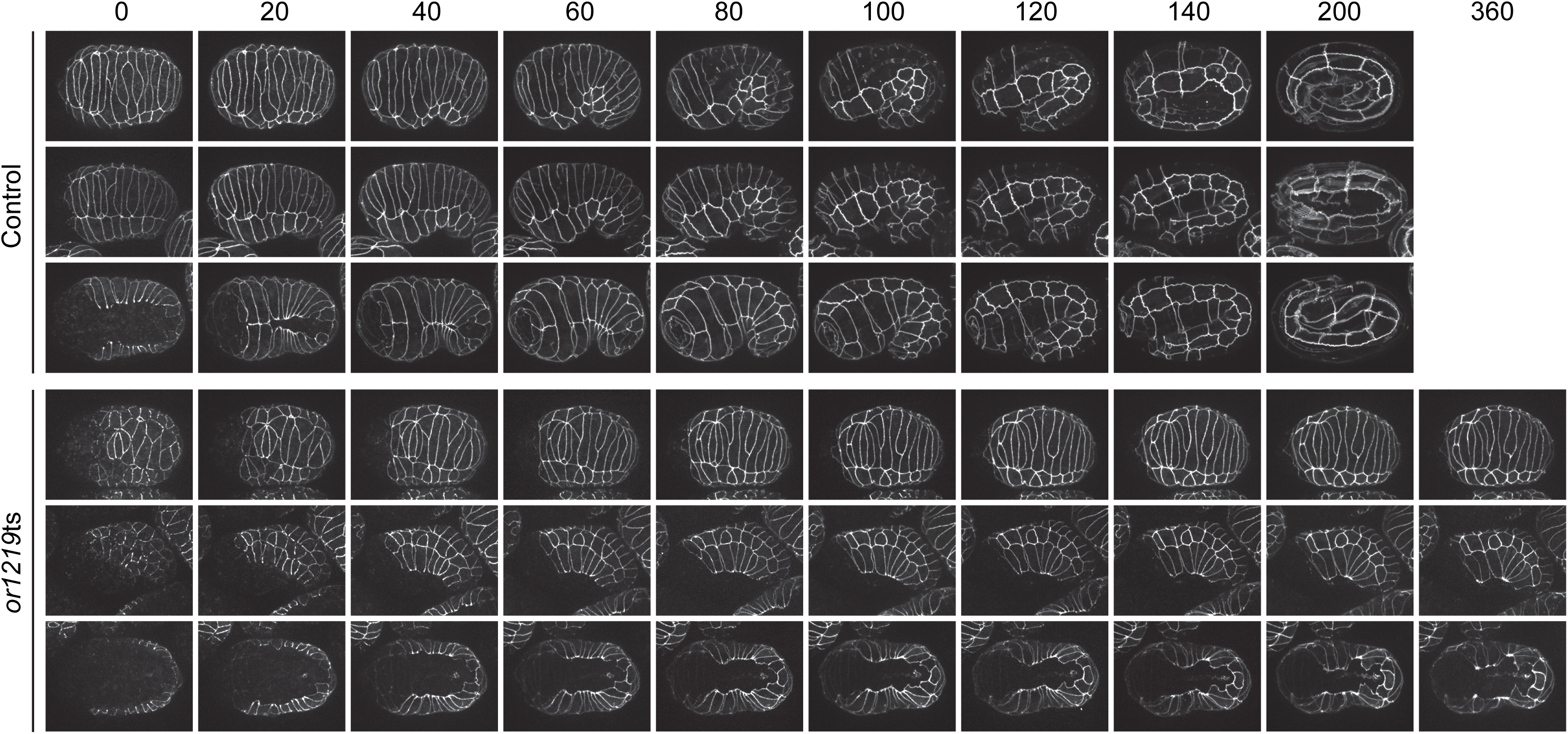
*or1219*ts embryos exhibit cell elongation failures after late upshifts. Maximum projection images of live control (top 3 rows) or mutant (*or1219*ts; bottom three rows) embryos, expressing DLG-1::GFP to mark epidermal cell membranes, after late upshifts. Minutes are listed across the top; time 0 corresponds to the start of imaging at the bean stage (one hour at 26°C after the late upshift time point; see Materials and Methods). Embryos were imaged every 20 minutes for 200 minutes (control embryos) or 360 minutes (*or1219*ts mutant embryos). Each row represents a single representative embryo over time in the dorsal orientation (top rows; 1 and 4), lateral orientation (middle rows; 2 and 5), and ventral orientation (bottom rows; 3 and 6). See text for a description of the defects in *or1219*ts mutants.

**Figure 5.**
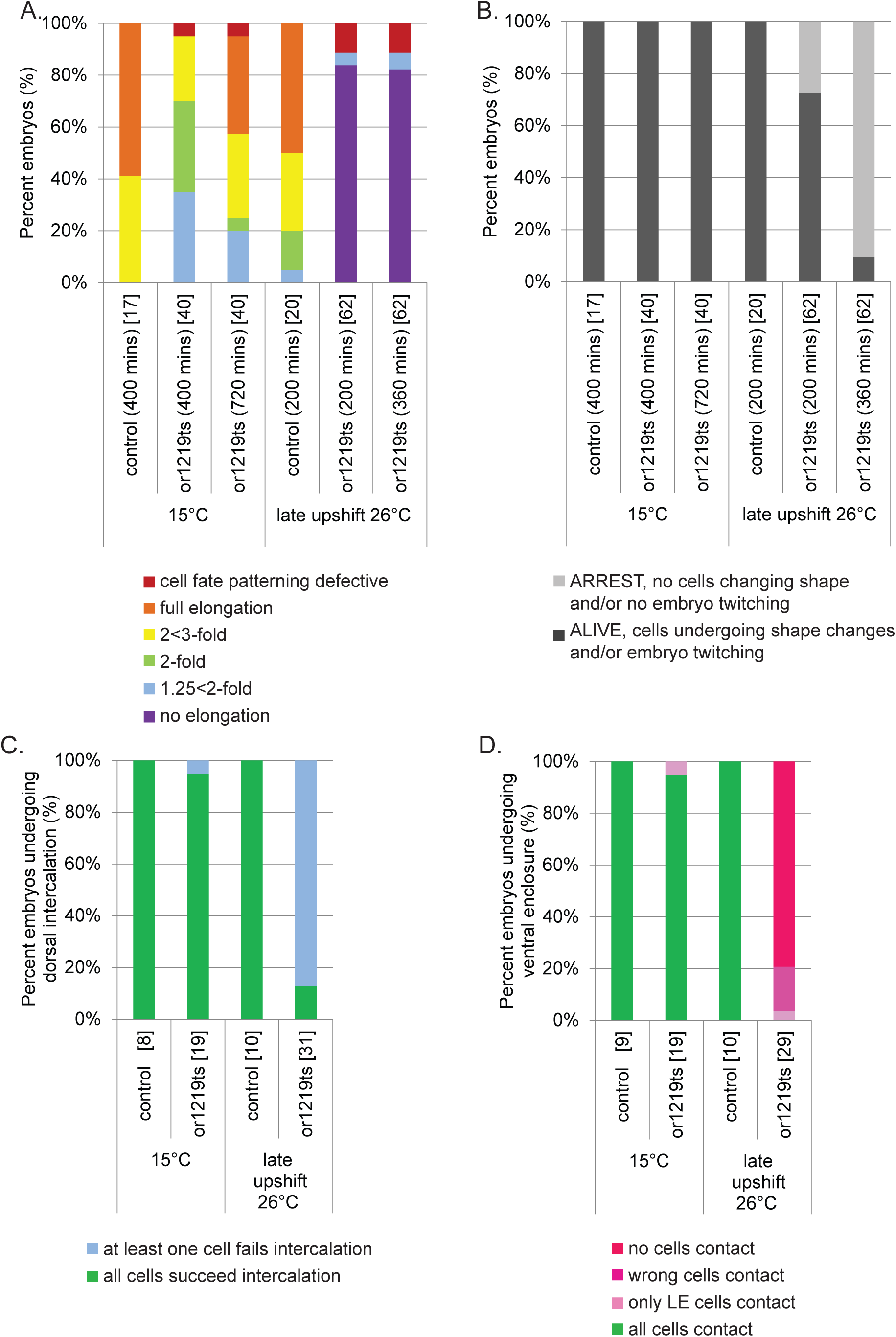
Quantification of the delayed development and elongation defects in *or1219*ts mutant embryos. **(A)** Quantification of terminal elongation-defective phenotypes for control and *or1219*ts embryos after live imaging for 400 or 720 minutes at 15°C, or for 200 or 360 minutes after late upshift, respectively. Percent of embryos that arrested with no elongation (purple), at 1.25<2-fold (light blue), at 2-fold (green), at 2<3-fold (yellow), full elongation (≥3-fold), or cell fate patterning defects (red) were scored using DLG-1::GFP to mark epidermal cell membranes and to view the body shape of the embryo. **(B)** Quantification of extent of viability for control and *or1219*ts embryos after live imaging for 400 or 720 minutes at 15°C or for 200 or 360 minutes after late upshift. Embryonic viability was assessed by comparing time points for each individual embryo and scoring cells undergoing shape changes and/or the whole embryo twitching. **(C)** Quantification of the extent of dorsal intercalation that occurred for control and *or1219*ts embryos by the final imaging time point at 15°C (400 or 720 minutes) and after late upshift (200 or 360 minutes). Embryos were scored by all dorsal epidermal cells succeeding at intercalation (extending across the dorsal midline and contacting the seam cells opposite to their starting point; green) or by one or more cells failing intercalation (failing to extend across the dorsal midline and make contact with the seam cells opposite to their starting point; light blue). (D) Quantification of the extent of ventral enclosure that occurred for control and *or1219*ts embryos by the final imaging time point at 15°C (400 or 720 minutes) and after late upshift (200 or 360 minutes). Embryos were scored for normal ventral enclosure (all ventral cells reaching the ventral midline; green), only the leading cells making contact but the ventral pocket remaining open (light pink), the wrong ventral cells making contact (non-leading cells; medium pink), or no ventral cells making contact (dark pink). The number of embryos scored for controls and mutants are in brackets.

We next examined *or1219*ts mutant embryos to determine if subsequent defects in epidermal cell shape or position after the late upshifts are associated with their failure to elongate. In control animals maintained at 15°C or upshifted to 26°C, dorsal interaction began when the two rows of 10 dorsal epidermal cells each elongated towards the dorsal midline. The dorsal cells then slid past one another to make contact with seam cells on the opposite side to produce a single “ladder-like” row of dorsal cells (Figures 3, 4). This morphogenetic event elongates the dorsal surface to create the embryo’s “bean-like” shape (Heid et al., 2001; Williams-Masson et al., 1998). In contrast to control embryos, in most *or1219*ts mutants after late temperature upshifts, one or more dorsal epidermal cells failed to make contact by 360 minutes with the seam cells on the opposite side (Figures 4, 5C). Furthermore, ventral enclosure also was perturbed in *or1219*ts mutants after the temperature upshift. In control animals, the ventral leading cells (the two most anteriorly located cells on either side of the ventral midline; cells V0, V1, V18, and V19) extended towards each other and made contact (Figures 3, 4, 5D). In most upshifted *or1219*ts mutants, the leading cells began to extend towards one another but ultimately failed and eventually retracted away from the ventral midline (Figures 4, 5D). In a few cases, the wrong ventral cells attempted to make contact across the ventral midline, or the leading cells made contact but the remaining posteriorly located ventral cells failed to enclose (Figures 5D, S9). Finally, we also observed that seam cells failed to elongate along the anterior-posterior axis (Figure 4). These results indicate that the *or1219*ts mutant is defective in epidermal morphogenesis and further support our conclusion that late stage upshifts are more likely to identify mutants specifically defective in embryonic morphogenesis.

## DISCUSSION

To identify genes required for embryonic morphogenesis in *C. elegans*, we have used a forward genetics approach that is biased only by a selection for conditional mutations in essential genes that lead to embryonic lethality at the restrictive temperature (TS-EL mutants). We initially examined the terminally differentiated embryonic phenotypes from nearly 200 TS-EL mutants that had normal early embryonic cell divisions. After upshifting L4 mutant larvae to the restrictive temperature, we found about 80 mutants that produced embryos with roughly normal cell fate patterning but nevertheless failed to fully elongate. Given the large proportion of mutants that failed to elongate, we suspected that low-penetrance or late defects in cell division or cell fate specification might indirectly result in defective elongation. Indeed, when we identified the causal mutations from a subset of these L4 upshifted mutants, most of the affected genes encode factors that function very generally in gene expression (see Table S5). We therefore refined our screen by upshifting embryos to the restrictive temperature just prior to the initiation of elongation, seeking to bypass earlier requirements and focus only on mutants that failed to elongate after late temperature upshifts. This reduced the number to 17 mutants with penetrant terminally differentiated elongation-defective phenotypes, and we have identified the likely causal mutations for eight of these mutants. Three of the affected genes—*rib-1*, *rib-2*, and *emb-9—*are known to be required for morphogenesis, indicating that our screening procedure was effective in identifying morphogenetic factors. Furthermore, one mutant, *or1219*ts, has defects in cell elongation during dorsal intercalation, ventral enclosure and seam cell elongation, cell shape changes that help drive embryonic elongation, further supporting the value of this mutant collection for advancing our understanding of the pathways that regulate and execute morphogenesis during *C. elegans* embryogenesis.

### The extracellular matrix and morphogenesis

The three genes *rib-1*, *rib-2*, and *emb-9* encode proteins that help form the extracellular matrix (ECM), which is known to influence morphogenetic processes in both invertebrates and vertebrates. The ECM is composed of fibrous structural proteins, including collagens and laminins, and proteoglycans, proteins that are covalently modified with glycosaminoglycan sugar chains including chondroitin-sulfate, dermatin-sulfate, keratin-sulfate, and heparan-sulfate (HS) (Clause and Barker, 2013; Daley and Yamada, 2013; Rozario and DeSimone, 2010).

#### rib-1 and rib-2 encode glycosyltransferases that affect morphogenetic signaling pathways

Our findings that *rib-1* and *rib-2* mutants have early defects in elongation are consistent with previous studies indicating that these enzymes play important roles during embryonic morphogenesis. The genes *rib-1* and *rib-2* both encode proteins involved in heparan-sulfate proteoglycan (HSPG) biosynthesis. RIB-2 is orthologous to human exostosin-like glycosyltransferase 3 (EXTL3) that catalyzes the addition of the first *N*-acetylglucosamine (GlcNAc) molecule at serine residues, while RIB-1 is orthologous to human exostosin glycosyltransferase 2 (EXT2) that elongates the linear HS chain by subsequently adding more GlcNAc molecules (Franks et al., 2006; Kitagawa et al., 2001; Kitagawa et al., 2007; Morio et al., 2003). Mutants of both *rib-1* and *rib-2* produce decreased levels of HS compared to wild-type worms, suggesting their functions are conserved for HSPG biosynthesis (Franks et al., 2006; Kitagawa et al., 2001; Kitagawa et al., 2007; Morio et al., 2003).

Glycosylation of ECM proteins has been linked to morphogenetic programs in other model organisms. HSPGs are composed of non-branched HS chains, alternating glucoronic acid and GlcNAc residues that are covalently linked to a core protein at serine amino acids (Poulain and Yost, 2015). HS chains can be attached to proteins anchored to the cell surface, either through transmembrane or by glycosylphophatidylinosiotol anchors, such as syndecans and glypicans, respectively. They can also be attached to secreted proteins in the extracellular space, including agrin, perlecan, and collagens IV and XVIII (Basak et al., 2016; Poulain and Yost, 2015). Due to their location near cell surfaces, HSPGs have been implicated in modulating signal transduction pathways that are important for cell survival, division, differentiation, and movement, in both invertebrates and vertebrates (Humphreys et al., 2013; Moulton et al., 2020; Poulain and Yost, 2015). Therefore, the “sugar code” hypothesis has been proposed (Holt and Dickson, 2005), whereby HS modifications to core proteins in specific tissues at specific times within the embryo control signaling pathways and coordinate morphogenesis and development. Although post-translational GlcNAc modifications to proteins are linked to regulating morphogenesis, the mechanisms by which they regulate cell movement and migration are not well understood. It is possible that HSPG’s function to help establish morphogen gradients by controlling ligand diffusion. HS chains may control levels of signaling by (1) sequestering the ligands in the ECM and thereby preventing their binding to receptors, (2) capturing ligands in the ECM to traffic them to their receptors, or (3) using transmembrane bound HSPGs to bring ligands close to their receptors (Poulain and Yost, 2015).

In *C. elegans*, *rib-1* and *rib-2* mutants have been previously reported to have maternal effect morphogenesis defects in embryonic elongation of both the pharynx and the whole body (Franks et al., 2006; Kitagawa et al., 2007; Morio et al., 2003), and microfilaments are disorganized in the pharynx of *rib-1* and *rib-2* mutants (Franks et al., 2006). These data suggest that *rib-1* and *rib-*2 are required for morphogenetic events, but how they influence epidermal cell shape changes or movements during elongation are unknown. *or1193ts* and *or1688*ts are the only known TS alleles for the genes *rib-1* and *rib-2*, respectively. Their penetrant elongation defects after late embryonic upshifts more clearly implicate HSPGs in embryonic elongation and provide useful tools for further investigation of their morphogenetic requirements. For example, temperature upshifts and downshifts at different times during embryogenesis may determine the critical time periods for *rib* gene functions. In addition, determining the spatial and temporal localization of both RIB proteins by making fluorescent protein fusions at endogenous loci using CRISPR/CAS-9 also may also help in understanding their morphogenetic requirements.

#### The basement membrane and body wall muscles are required for later stages of elongation

Our identification of a mutation in *hlh-1* is consistent with the BM being required for late stages of elongation in response to body wall muscle contractions. Mutants deficient in muscle development result in the “Pat” phenotype, <u>p</u>aralyzed <u>a</u>rrest at two-fold, because a mechanotransduction pathway requires muscle contractions for elongation past the 2- fold stage, with muscles attached to the epidermis via BM components (Labouesse, 2012; Zhang et al., 2011). Our screen identified *hlh-1,* which encodes a bHLH transcription factor conserved with human MyoD that has roles in muscle differentiation. HLH-1 is sufficient, but not required, to specify body wall muscles during embryogenesis and regulates expression of muscle chaperone proteins during muscle differentiation. Loss of *hlh-1* has been shown previously to result in arrest at the 2-fold stage of elongation, with dis-organized body wall muscles that have reduced contractions (Bar-Lavan et al., 2016; Chen et al., 1992; Chen et al., 1994; Fukushige and Krause, 2005; Harfe et al., 1998; Krause et al., 1990).

EMB-9 is orthologous to the human type IV collagen α1 subunit (COL4A5), which localizes to the basal side of the basement membrane (BM) (Guo and Kramer, 1989). The BM is composed of two polymeric networks, laminin and collagen, that are thought to be joined by bridging molecules such as nidogen and perlecan (Jayadev and Sherwood, 2017). Laminin interacts with cell surfaces by binding adhesion receptors and lipids, while less is known about whether collagen directly associates with epithelial cells (Jayadev and Sherwood, 2017). Collagen is thought to provide tensile strength to the BM, protecting it from mechanical stresses, preventing tissue distensibility, and sterically limiting tissue expansion, thereby affecting epithelial morphogenesis. For example, collagen IV is required for the rotation and subsequent elongation of *Drosophila* egg chambers (Haigo and Bilder, 2011), and collagen is known to restrict branching morphogenesis during mammary gland development (Fata et al., 2004; Hinck and Silberstein, 2005).

Our identification of a mutation in *emb-9* is consistent with previous studies indicating a role for BM during later stages of embryonic elongation. In *C. elegans*, EMB-9 is expressed in body wall muscle cells by the 1.25-fold (comma) stage and surrounds the body wall and pharyngeal muscle cells (Graham et al., 1997; Gupta et al., 1997; Keeley et al., 2020). The absence of collagen and other BM components are lethal in *C. elegans*, *Drosophila*, and mouse embryos. In *C. elegans*, loss of the collagen bridging molecule perlecan (*unc-52*) and both collagens (*emb-9* and *let-2*) lead to the Pat phenotype (Gupta et al., 1997; Hresko et al., 1994). Five previously isolated TS-EL mutations in *emb-9* result in embryos that produce normal patterns of cell fate but arrest shortly after muscle twitching has initiated, at either the 2-fold or 3-fold stage depending on the mutant allele (Gupta et al., 1997; Miwa et al., 1980; Schierenberg et al., 1980). The *emb-9(or1723*ts) allele that we have isolated introduces a unique missense mutation compared to the five existing TS-EL mutations and similarly arrests at the 2- fold stage or slightly later, when muscle twitching begins. In *C. elegans* and *Drosophila*, muscle cells detach from the body wall during muscle contractions after depletion of collagen, suggesting the BM is unable to withstand mechanical stress (Jayadev and Sherwood, 2017). Again, our identification of a TS allele of *emb-9*, and the penetrant elongation defects we observed after late temperature upshifts, more clearly implicate *emb-9* in *C. elegans* embryonic morphogenesis.

### Cell shape defects in *or1219*ts mutants

In addition to identifying causal mutations in genes that have been previously implicated in morphogenesis, we also have characterized the epidermal cell shape defects in another mutant, *or1219*ts, using live imaging and DLG-1::GFP, a transgenically expressed fluorescent protein fusion that marks epidermal cell membranes. Although the causal mutation has yet to be identified, we chose to examine epidermal cell shape changes in *or1219*ts because it had the most penetrant, fully elongation-defective phenotype after late upshifts. In most *or1219*ts embryos after late upshifts, one or more dorsal epidermal cells failed to fully extend across the dorsal surface to make contact with seam cells on the opposite side. The pair of ventral leading cells on either side of the embryo also displayed elongation defects, failing to extend towards the ventral midline or to stably adhere to one another. In addition, the subsequent closure of the remaining posterior pocket cells was unsuccessful. Since seam cell elongation requires a complete epidermal sheet, as shown by laser ablation (Priess and Hirsh, 1986), the failure of seam cells to elongate along the anterior-posterior axis in *or1219*ts mutants might be indirectly due to the earlier defects in dorsal intercalation and ventral enclosure. These defects in *or1219*ts epidermal cell shape changes during embryonic elongation further indicate that our screen has identified mutants specifically required for epidermal morphogenesis.

### Novel gene requirements during morphogenesis

The possible morphogenetic roles played by the remaining three genes we have identified in our screen are less obvious. *zim-3* encodes a protein containing two zinc-finger domains that is thus far known to be conserved only in nematodes. ZIM-3 is required for homologous pairing of the autosomal chromosomes I and IV during meiosis and interacts with the nuclear lamina (Phillips and Dernburg, 2006; Phillips et al., 2009). *emb-4* encodes a nuclear protein with a putative AAA ATPase domain similar to helicases of the DEAD-box family and is orthologous to mammalian Aquarius, that functions in pre-mRNA splicing (De et al., 2015; Katic and Greenwald, 2006; Wahl and Luhrmann, 2015). EMB-4 has also been implicated in transcriptional activation and cell fate specification. Loss of *emb-4* results in slowed E-cell lineage cell division (Schierenberg et al., 1980), in embryos that develop dis-organized tissues (Checchi and Kelly, 2006), in defective chromatin remodeling in the germline precursor cells Z2/Z3 (Checchi and Kelly, 2006), and in intron-dependent transcriptional gene silencing by RNAi (Akay et al., 2017). EMB-4 functions as a nonessential positive regulator of Notch signaling (Checchi and Kelly, 2006; Katic and Greenwald, 2006), but the exact biochemical role of EMB-4 is still unknown. Lastly, *emb-5* encodes a RNA polymerase II transcription elongation factor orthologous to human SPT6, which interacts with histones and likely functions to displace nucleosomes during transcription (Kwak and Lis, 2013; Nishiwaki et al., 1993). Like EMB-4, EMB-5 has a role in the timing of E-cell lineage cell divisions during gastrulation and is likely a positive regulator downstream of the Notch signaling pathway (Hubbard et al., 1996; Miwa et al., 1980; Nishiwaki et al., 1993). Three of these genes—*zim-3*, *emb-4*, and *emb-5*—all have known early requirements during development. However, our results indicate that they also have previously uncharacterized mid to late embryonic requirements since they all fail to properly elongate after late upshifts, while elongating more normally if kept at the permissive temperature. Because *zim-3*, *emb-4*, and *emb-5* all lack a relationship to known requirements to morphogenesis in *C. elegans* and other systems, they may represent new and interesting morphogenetic factors.

Thus far our screening efforts have not identified cytoskeletal regulators that have been previously shown to influence morphogenesis. This could be due to a lack of saturation in screening for relatively rare conditional mutations. In addition, genes and their encoded proteins likely vary with respect to how susceptible they are to acquiring mutations that confer conditional, temperature-dependent function.

### Future directions for advancing our understanding of *C. elegans* embryonic morphogenesis

Our screen provides a foundation for further investigation of the mechanisms that regulate and execute *C. elegans* embryonic morphogenesis. The causal mutations for nine alleles remain to be identified, and it may be necessary to generate deletion alleles using CRISPR for complementation tests, as alleles for several candidate genes do not exist. Furthermore, it will be important to confirm the identity of all causal mutations by using CRISPR technology to recreate the putative causal mutations, or by using transgenic rescue with the wild-type candidate genes. Live imaging to examine epidermal cell shape dynamics during morphogenesis in the remaining 16 penetrant morphogenesis-defective late-upshift mutants, and more detailed analysis of cell fate specification and cell division in these mutants (Bao et al., 2006; Moore et al., 2013; Murray et al., 2008; Santella et al., 2010), should also prove informative. Finally, further unbiased forward genetic screens, focusing on either existing or newly isolated mutants after chemical mutagenesis, or taking advantage of a new eggshell-permeable analog of auxin to pursue more systematic screens using auxin-inducible degradation (Negishi et al., 2019), may identify important new morphogenetic factors.

## ACKNOWLEDGMENTS

We thank past members of the Bowerman lab in helping to isolate conditional mutants, Chris Doe and Diana Libuda for sharing laboratory equipment, Adam Fries from the University of Oregon Imaging Core Facility for microscopy advice, Doug Turnbull and the University of Oregon Genomics and Cell Characterization Core Facility (GC3F) for Illumina sequencing and bioinformatics expertise, members of the Bowerman, Libuda and Andrew Chisholm laboratories for helpful discussions, Cherry Biotech for equipment support, the Caenorhabditis Genetics Center (CGC; University of Minnesota, Twin Cities, Minnesota which is funded by NIH Office of Research Infrastructure Programs; P40 OD010440) and the National Bioresource Project (NBRP; Department of Physiology, Tokyo Women’s Medical University School of Medicine, Tokyo, Japan) for *C. elegans* strains.

## COMPETING INTERESTS

The authors declare no competing or financial interests.

## AUTHOR CONTRIBUTIONS

Conceptualization: MCJ, JL, ZB, BB; Methodology: MCJ, JL, TP, EC, YY, FF, AM, DH, AH, MAZ, HS, NT; Analysis: MCJ, JL; Data curation: MCJ, JL, BB; Writing: MCJ, BB; Supervision: BB; Project administration: BB; Funding acquisition: MCJ, BB, and ZB.

## FUNDING

This work was supported by the National Institutes of Health [GM114053 to BB and ZB; GM049869 and GM131749 to BB; GM126677 to MCJ; R25-HD070817 to Judith Eisen that supported AM and FF in the Oregon Undergraduate Researchers in SPUR; P30 CA008748 to MSKCC for Cancer Center Support/Core], the National Science Foundation [DBI 1758015 to Peter O’Day and Alice Barkan that supported MAZ in the University of Oregon Summer Program for Undergraduate Research (SPUR)], and the McNair Scholars Program [supported TP].

## SUPPLEMENTAL FIGURE LEGENDS

**Supplemental Figure 1.**
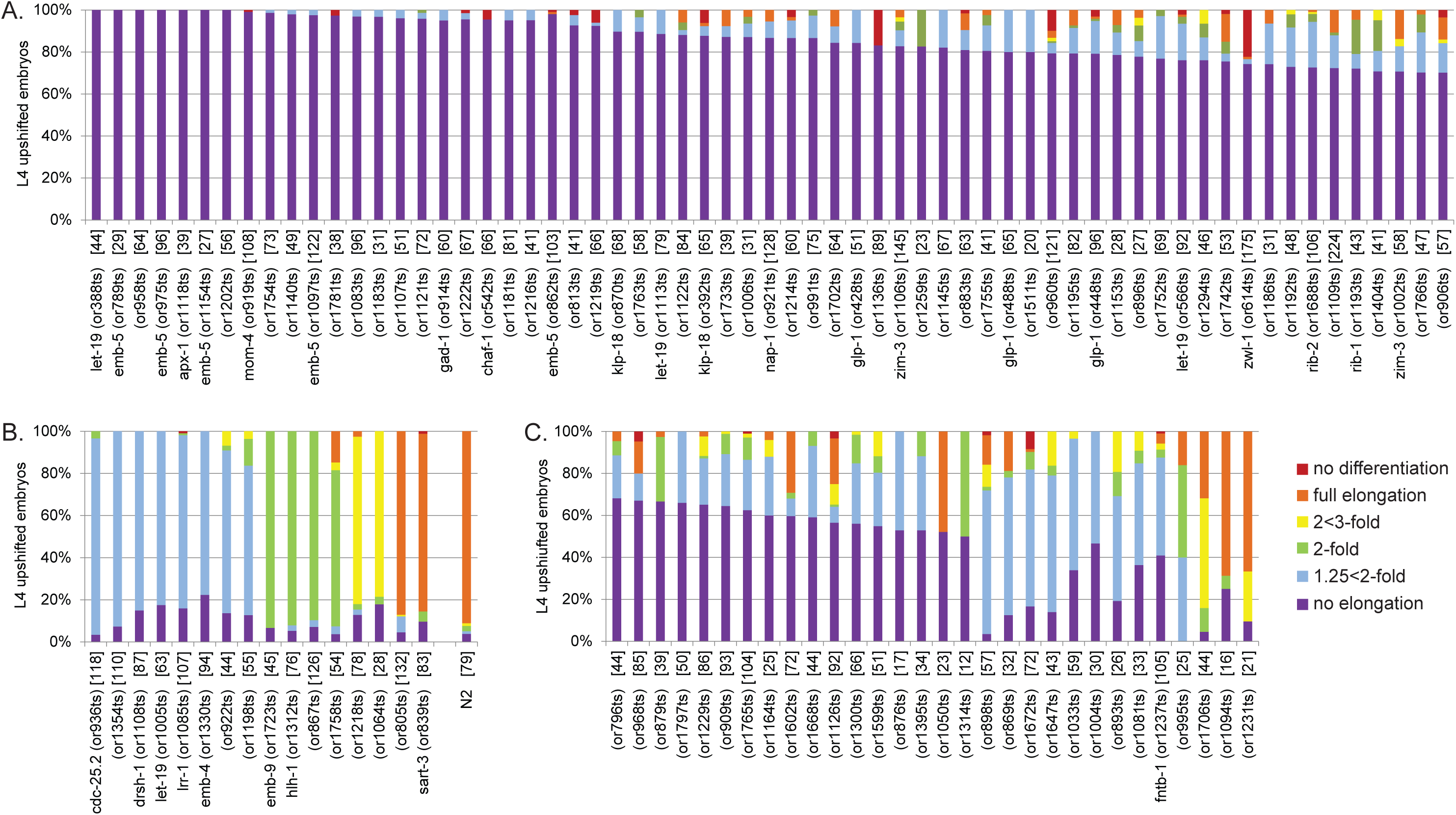
Penetrant and variable terminal elongation-defective phenotypes after L4 upshifts for TS-EL mutants. **(A-B)** Quantification of 79 penetrant terminal elongation-defective phenotypes for TS-EL mutant (≥70% embryos arrested with similar extents of elongation) and wild-type (N2) embryos after L4 upshifts. 63 TS-EL mutants arrested with 70% or more exhibiting no elongation (A). 16 TS-EL mutants arrested with 70% or more exhibiting similar less extensive elongation arrests (B). **(C)** Quantification of 30 variable terminal elongation-defective phenotypes for TS-EL mutant embryos after L4 upshift (40-70% embryos arrested with a similar extent of elongation). Percent of embryos that differentiated well but arrested with no elongation (purple), or arrested at the 1.25<2-fold stage (light blue), at the 2-fold stage (green), at the 2<3-fold stage (yellow), or with full elongation (≥3-fold), or exhibited differentiation defects (red) were scored using Nomarski optics. Examples of Nomarski images for control embryos at each elongation stage are shown in the Key in Figure 1. TS-EL mutants are shown from left to right by decreasing penetrance for the most penetrant elongation category. If the causal mutation has been identified for a TS-EL mutant, the affected gene is listed next to the TS-EL mutant allele, and the number of embryos scored for each mutant is in brackets.

**Supplemental Figure 2.**
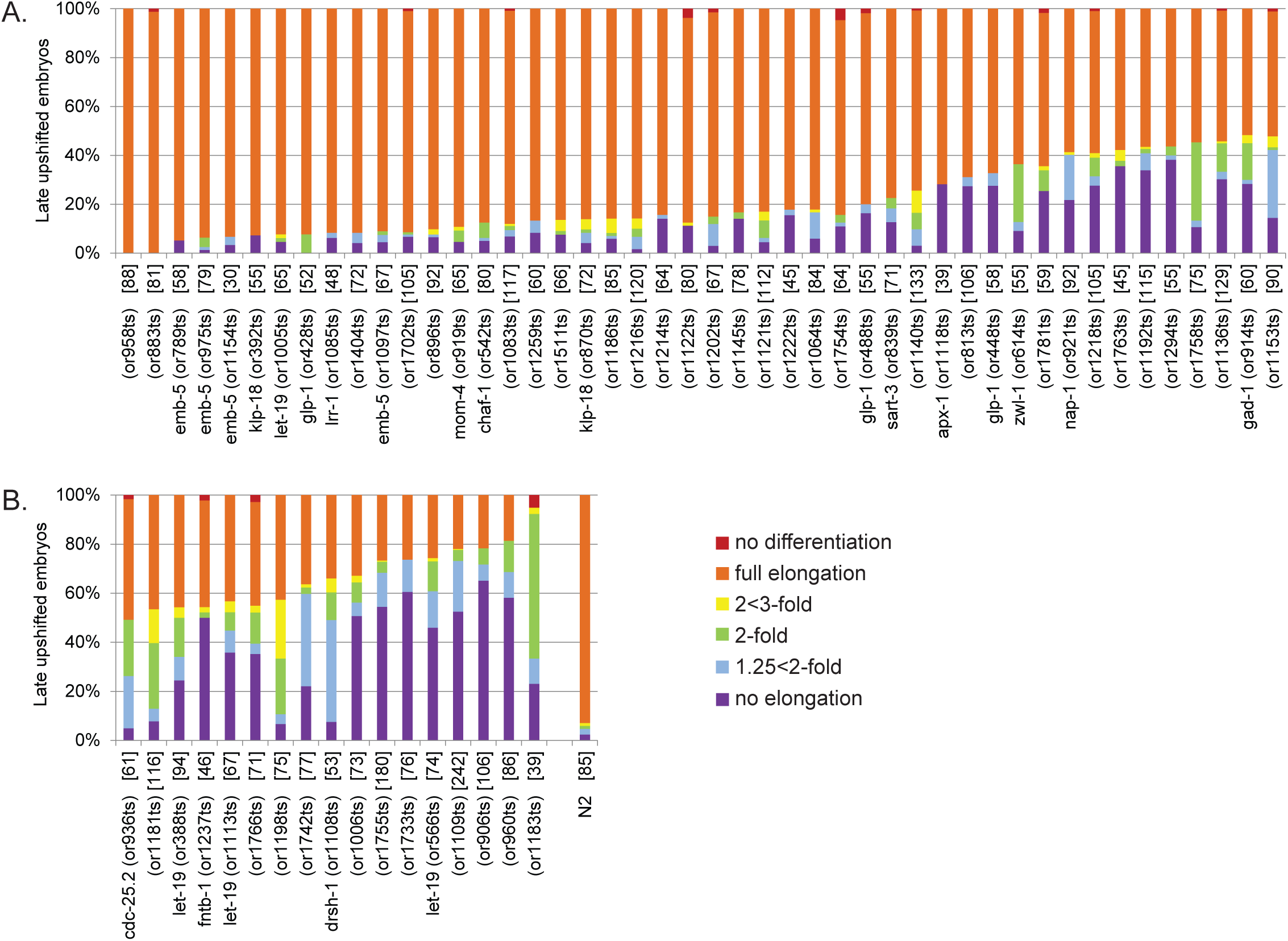
Quantification of less penetrant elongation defects in TS-EL mutants after late upshifts. Quantification of terminal elongation-defective phenotypes for TS-EL mutant and wild-type (N2) embryos after late upshifts. **(A)** TS-EL mutants with penetrant elongation defects after L4 upshifts that exhibited weak defects after late upshifts. 50-100% of embryos undergo full elongation and hatch and/or twitch. **(B)** TS-EL mutants with penetrant elongation defects after L4 upshifts that exhibited variable elongation defects after late upshifts. Percent of embryos that differentiated well but arrested with no elongation (purple), or arrested at the 1.25<2-fold stage (light blue), at the 2-fold stage (green), at the 2<3-fold stage (yellow), or with full elongation (≥3-fold), or exhibited differentiation defects (red) were scored using Nomarski optics. Examples of Nomarski images for control embryos at each elongation stage are shown in the Key in Figure 1. TS-EL mutants are shown from left to right by decreasing penetrance for the full elongation category. If the causal mutation has been identified for a TS-EL mutant, the affected gene is listed next to the TS-EL mutant allele, and the number of embryos scored for each mutant is in brackets.

**Supplemental Figure 3.**
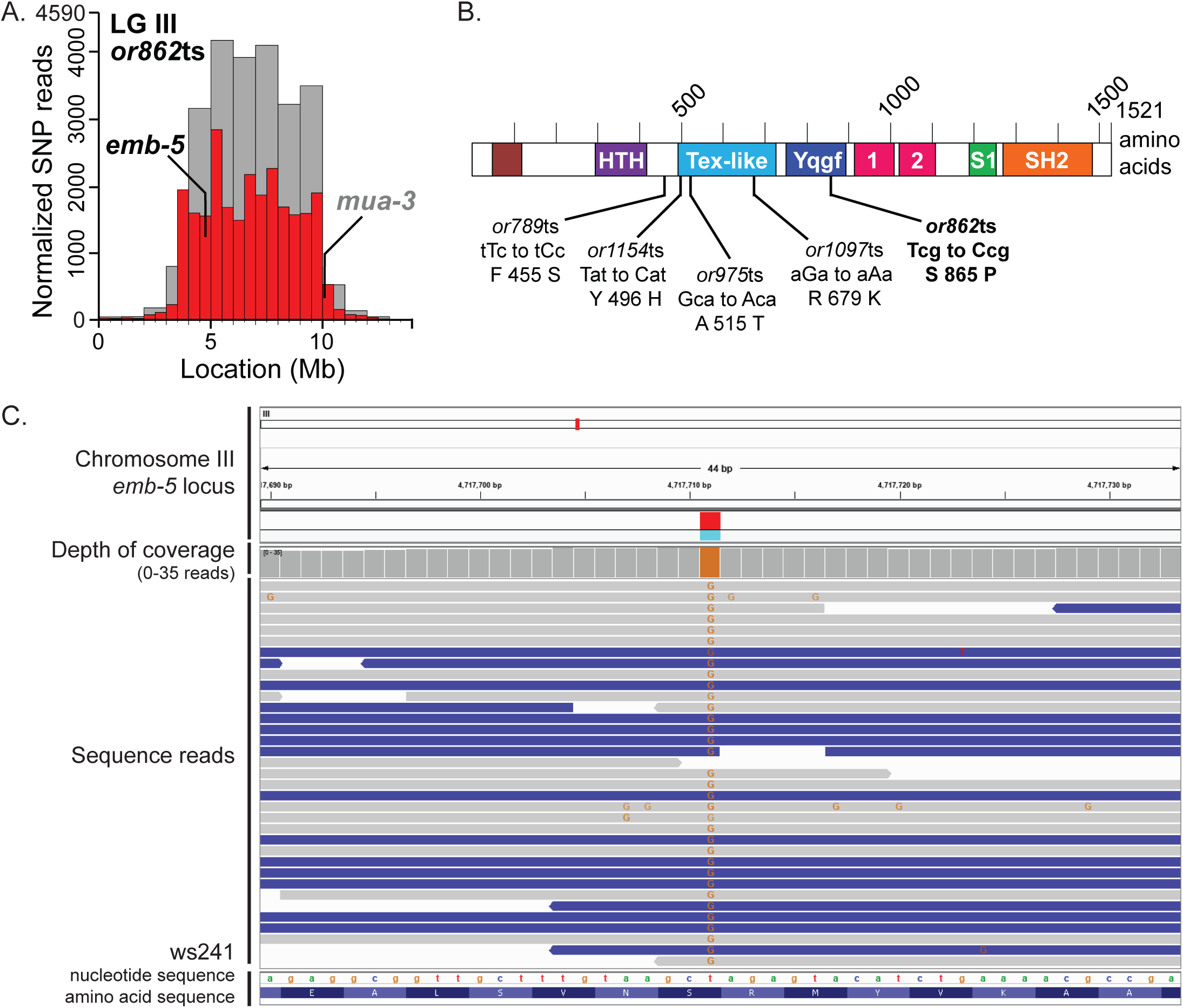
Identification of putative *emb-5(or862*ts) causal mutation. **(A)** SNP mapping data for the *or862*ts mutant on linkage group III with identified causal mutations, showing the frequency of homozygous parental alleles plotted against chromosomal position in bins of either 1 megabase (gray bars) or 0.5 megabase (red bars). Candidate essential genes in which a missense mutation (*emb-5*) or splice site mutation (*mua-3*) were detected are indicated. For complementation test in which *or862*ts failed to complement the known genetic mutation, the gene is dark and bolded; for complementation test in which *or862*ts complemented the known mutation, the gene is gray and bolded. See Table 2 for complementation test results. **(B)** Sequence alterations in the predicted EMB-5 protein for *or862*ts and the other 4 alleles identified by our screen (*or789*ts, *or975*ts, *or1097*ts, and *or1154*ts). The causal mutation for *hc61*ts, the mutant allele used for the complementation test, has not been curated. EMB-5 is orthologous to human SPT6, a transcription elongation factor (Kwak and Lis, 2013; Nishiwaki et al., 1993). EMB-5 contains an acidic N-terminal domain (brown), a helix-turn-helix DNA binding domain (HTH, purple), a Tex-like domain (light blue), an RNase H-like domain (Yqgf, dark blue), 2 RuvA 2-like domains (pink), an S1 RNA binding domain (green) and an SH2 domain (orange; INTERPRO and Pfam on https://wormbase.org). **(C)** Integrative Genomics Viewer (Broad Institute) screenshot of the sequencing reads at the site of the *or862*ts missense mutation at the *emb-5* locus. The orange bar in the depth of coverage section indicates the homozygosity of the T to C nucleotide change across the reads. Gray lines indicate all bases matched the reference sequence; blue lines imply reads of the opposite strand (https://software.broadinstitute.org/software/igv/interpreting_pair_orientations). Single nucleotide changes are indicated on each read (green A, blue C, orange G, and red T). Nucleotide and amino acid sequences are read from right to left.

**Supplemental Figure 4.**
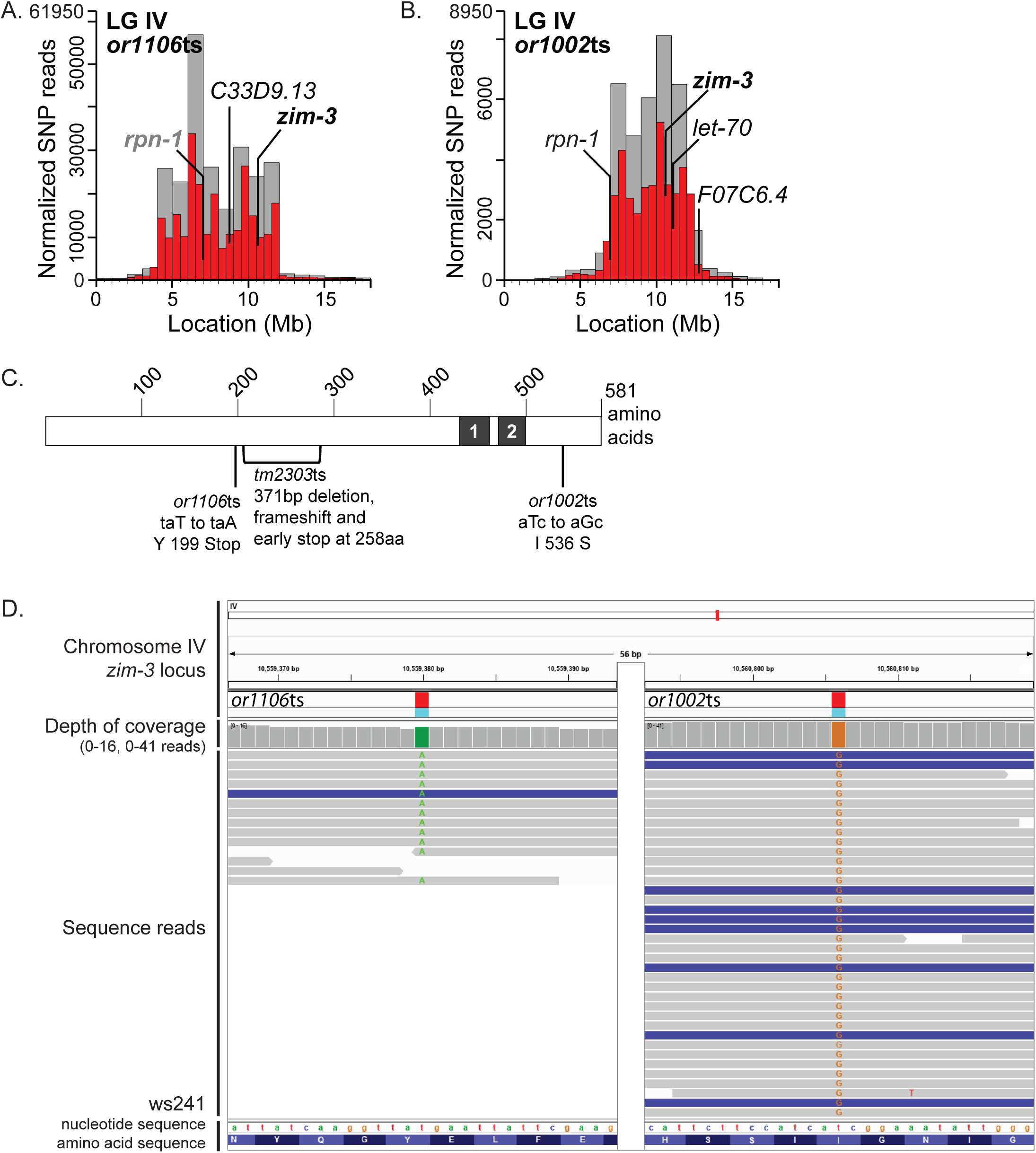
Identification of putative *zim-3*(*or1106*ts and *or1002*ts) causal mutations. **(A-B)** SNP mapping data for *or1106*ts (A) and *or1002*ts (B) mutants on linkage group IV with identified causal mutations, showing the frequency of homozygous parental alleles plotted against chromosomal position in bins of either 1 megabase (gray bars) or 0.5 megabase (red bars). Candidate essential genes in which missense mutations (*or1106*ts: *rpn-1* and *C33D9.13*; *or1002*ts: *zim-3*, *rpn-1*, *let-70*, and *F07C6.4*) or a nonsense mutation (*or1106*ts: *zim-3*) were detected are indicated. For complementation tests in which the TS-EL mutants failed to complement the known genetic mutation, the gene is dark and bolded; for complementation test in which the TS-EL mutant complemented the known mutation, the gene is gray and bolded; for genes in which a complementation test was not performed, genes are not bolded. See Table 2 for complementation test results. **(C)** Sequence alterations in the predicted ZIM-3 protein, isoform a, for *or1106*ts, *or1002*ts, and *tm2303*ts, the known mutant allele used for the complementation test. ZIM-3 is a *Caenorhabditis* specific gene required for homologous pairing of chromosomes during meiosis and contains 2 zinc-finger domains (black box) (Phillips and Dernburg, 2006; Phillips et al., 2009). **(D)** Integrative Genomics Viewer (Broad Institute) screenshot of the sequencing reads at the site of the *or1106*ts nonsense mutation (LEFT) and the *or1002*ts missense mutation (RIGHT) at the *zim-3* locus. The green bar in the depth of coverage section indicates the homozygosity of the T to A nucleotide change across the reads for *or1106*ts. The orange bar in the depth of coverage section indicates the homozygosity of the T to G nucleotide change across the reads for *or1002*ts. Gray lines indicate all bases matched the reference sequence; blue lines imply reads of the opposite strand (https://software.broadinstitute.org/software/igv/interpreting_pair_orientations). Single nucleotide changes are indicated on each read (green A, blue C, orange G, and red T). Nucleotide and amino acid sequences are read from left to right.

**Supplemental Figure 5.**
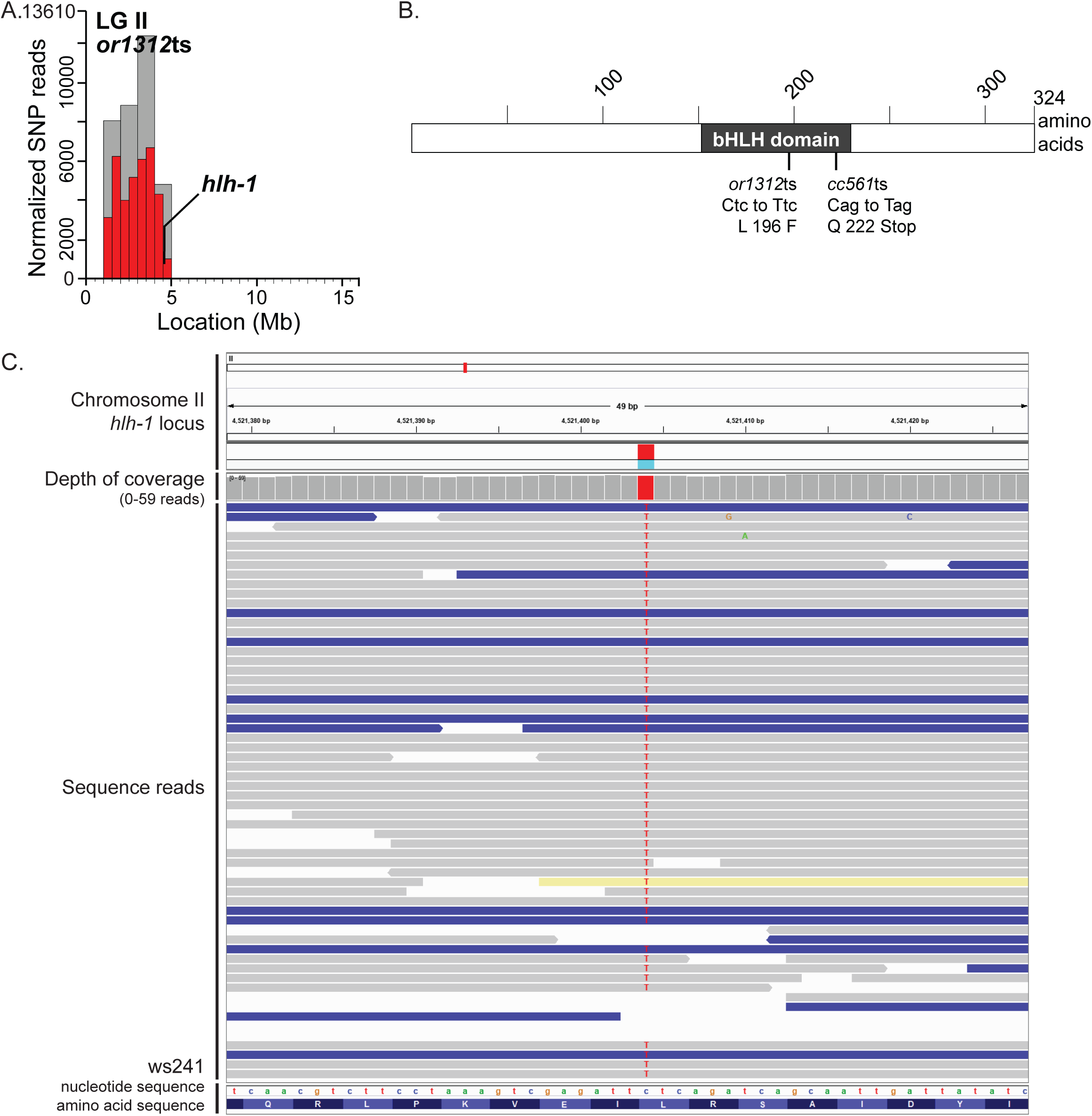
Identification of putative *hlh-1*(*or1312*ts) causal mutation. **(A)** SNP mapping data for the *or1312*ts mutant on linkage group II with identified causal mutations, showing the frequency of homozygous parental alleles plotted against chromosomal position in bins of either 1 megabase (gray bars) or 0.5 megabase (red bars). Candidate essential gene in which a missense mutation was detected is indicated. For complementation test in which *or1312*ts failed to complement the known genetic mutation, the gene is dark and bolded. See Table 2 for complementation test results. **(B)** Sequence alterations in the predicted HLH-1 protein, isoform b, for *or1312*ts and *cc561*ts, the known mutant allele used for the complementation test. HLH-1 is orthologous to human MyoD, a basic helix-loop-helix (bHLH) transcription factor (Krause et al., 1990), and contains a bHLH domain (black box corresponding to amino acids 155-231; INTERPRO, Pfam, smart, and superfamily on https://wormbase.org). **(C)** Integrative Genomics Viewer (Broad Institute) screenshot of the sequencing reads at the site of the *or1312*ts missense mutation at the *hlh-1* locus. The red bar in the depth of coverage section indicates the homozygosity of the C to T nucleotide change across the reads. Gray lines indicate all bases matched the reference sequence; blue lines imply reads of the opposite strand (https://software.broadinstitute.org/software/igv/interpreting_pair_orientations). The light yellow read implies the mate pair is on linkage group IV instead of II. Single nucleotide changes are indicated on each read (green A, blue C, orange G, and red T). Nucleotide and amino acid sequences are read from left to right.

**Supplemental Figure 6.**
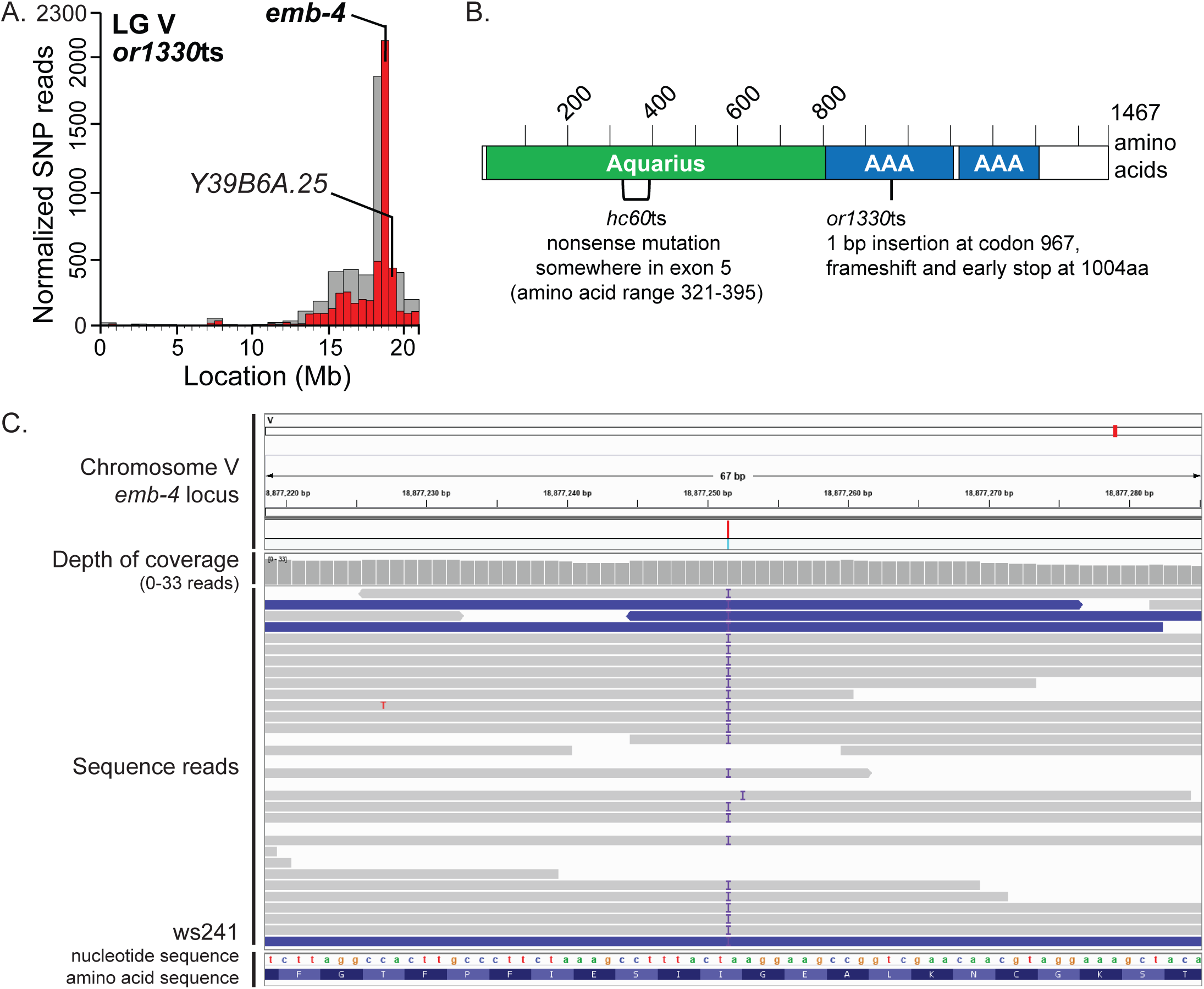
Identification of putative *emb-4*(*or1330*ts) causal mutation. **(A)** SNP mapping data for the *or1330*ts mutant on linkage group V with identified causal mutations, showing the frequency of homozygous parental alleles plotted against chromosomal position in bins of either 1 megabase (gray bars) or 0.5 megabase (red bars). Candidate essential genes in which a missense mutation (*Y39B6A.25*) or a 1 bp insertion (*emb-4*) were detected are indicated. For complementation test in which *or1330*ts failed to complement the known genetic mutation, the gene is dark and bolded; for the gene in which a complementation test was not performed, the gene is not bolded. See Table 2 for complementation test results. **(B)** Sequence alterations in the predicted EMB-4 protein, isoform a, for *or1330*ts and *hc60*ts, the known mutant allele used for the complementation test (Checchi and Kelly, 2006). EMB-4 is orthologous to human Aquarius (Katic and Greenwald, 2006) and contains an N-terminus Aquarius intron-binding domain (green) and 2 AAA ATPase DEAD-box helicase domains (blue; INTERPRO and Pfam on https://wormbase.org). **(C)** Integrative Genomics Viewer (Broad Institute) screenshot of the sequencing reads at the site of the *or1330*ts missense mutation at the *emb-4* locus. The purple “I” indicates an insertion of an A into codon 967. Gray lines indicate all bases matched the reference sequence; blue lines imply reads of the opposite strand (https://software.broadinstitute.org/software/igv/interpreting_pair_orientations). Single nucleotide changes are indicated on each read (green A, blue C, orange G, and red T). Nucleotide and amino acid sequences are read from right to left.

**Supplemental Figure 7.**
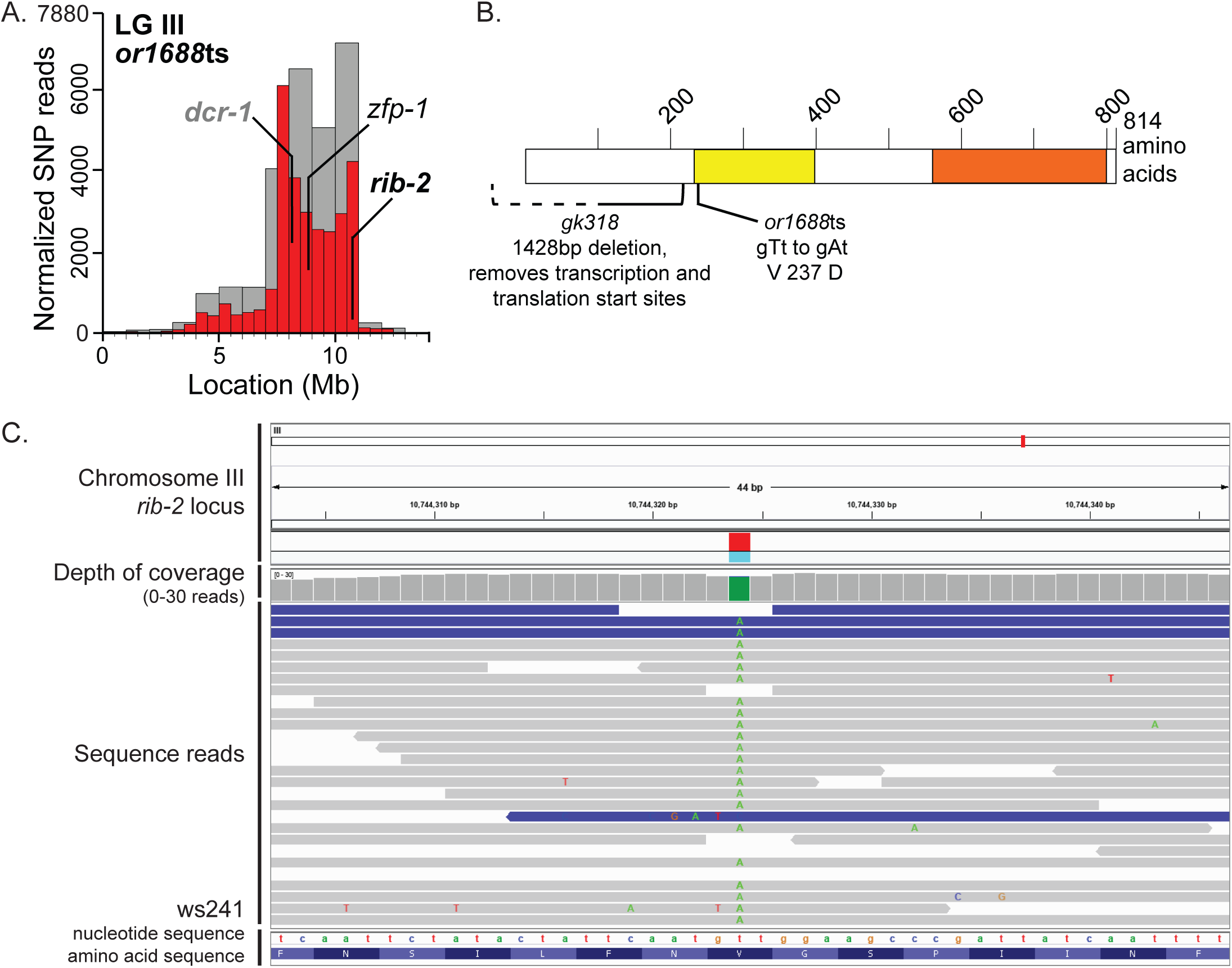
Identification of putative *rib-2*(*or1688*ts) causal mutation. **(A)** SNP mapping data for the *or1688*ts mutant on linkage group III with identified causal mutations, showing the frequency of homozygous parental alleles plotted against chromosomal position in bins of either 1 megabase (gray bars) or 0.5 megabase (red bars). Candidate essential genes in which missense mutations (*rib-2* and *zfp-1*) or splice site mutations (*dcr-1*) were detected are indicated. For complementation test in which *or1688*ts failed to complement the known genetic mutation, the gene is dark and bolded; for complementation test in which *or688*ts complemented the known mutation, the gene is gray and bolded; for the gene in which a complementation test was not performed, the gene is not bolded. See Table 2 for complementation test results. **(B)** Sequence alterations in the predicted RIB-2 protein for *or1688*ts and *gk318*, the known mutant allele used for the complementation test. RIB-2 is orthologous to human exostosin-like glycosyltransferase 3 (EXT3) (Kitagawa et al., 2007) and contains an exostosin-like domain (yellow) and a glycosyltransferase domain (orange; hmmpanther, INTERPRO, Pfam, and superfamily on https://wormbase.org). **(C)** Integrative Genomics Viewer (Broad Institute) screenshot of the sequencing reads at the site of the *or1688*ts missense mutation at the *rib-2* locus. The green bar in the depth of coverage section indicates the homozygosity of the T to A nucleotide change across the reads. Gray lines indicate all bases matched the reference sequence; blue lines imply reads of the opposite strand (https://software.broadinstitute.org/software/igv/interpreting_pair_orientations). Single nucleotide changes are indicated on each read (green A, blue C, orange G, and red T). Nucleotide and amino acid sequences are read from left to right.

**Supplemental Figure 8.**
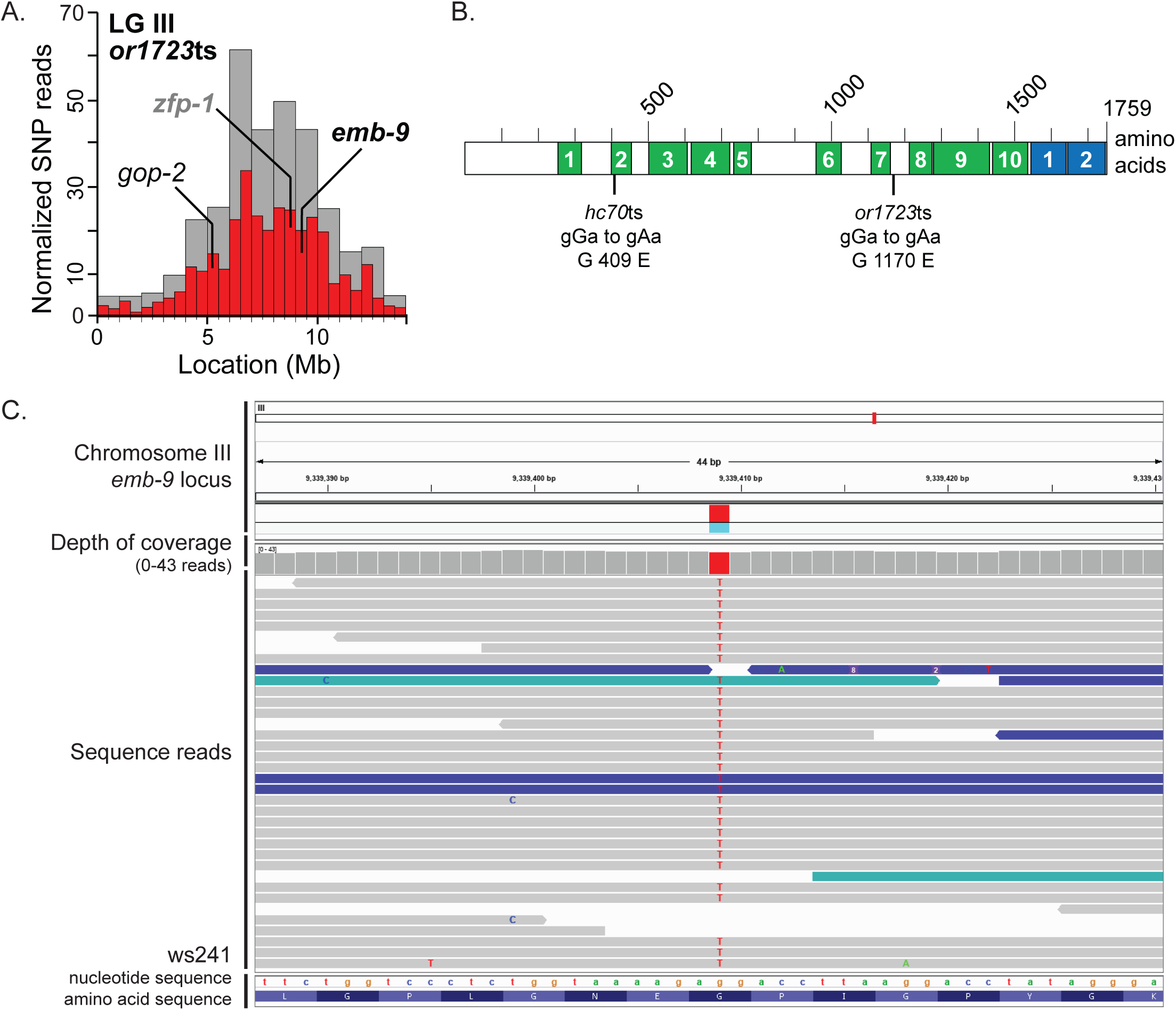
Identification of putative *emb-9* (*or1723*ts). **(A)** SNP mapping data for the *or1723*ts mutant on linkage group III with identified causal mutations, showing the frequency of homozygous parental alleles plotted against chromosomal position in bins of either 1 megabase (gray bars) or 0.5 megabase (red bars). Candidate essential genes in which missense mutations were detected are indicated. For complementation test in which *or1723*ts failed to complement the known genetic mutation, the gene is dark and bolded; for complementation test in which *or1723*ts complemented the known mutation, the gene is gray and bolded; for the gene in which a complementation test was not performed, the gene is not bolded. See Table 2 for complementation test results. **(B)** Sequence alterations in the predicted EMB-9 protein, isoform a, for *or1723*ts and *hc70*ts, the known mutant allele used for the complementation test. EMB-9 is orthologous to human type IV collagen α1 subunit (COL4A5) (Guo and Kramer, 1989). EMB-9 contains 10 collagen triple helix repeat domains (green) and 2 c-type lectin fold domains (blue; INTERPRO and Pfam on https://wormbase.org). **(C)** Integrative Genomics Viewer (Broad Institute) screenshot of the sequencing reads at the site of the *or1723*ts missense mutation at the *emb-9* locus. The red bar in the depth of coverage section indicates the homozygosity of the G to A nucleotide change across the reads. Gray lines indicate all bases matched the reference sequence; blue and teal lines imply reads of the opposite strand (https://software.broadinstitute.org/software/igv/interpreting_pair_orientations). Single nucleotide changes are indicated on each read (green A, blue C, orange G, and red T). Nucleotide and amino acid sequences are read from right to left.

**Supplemental Figure 9.**
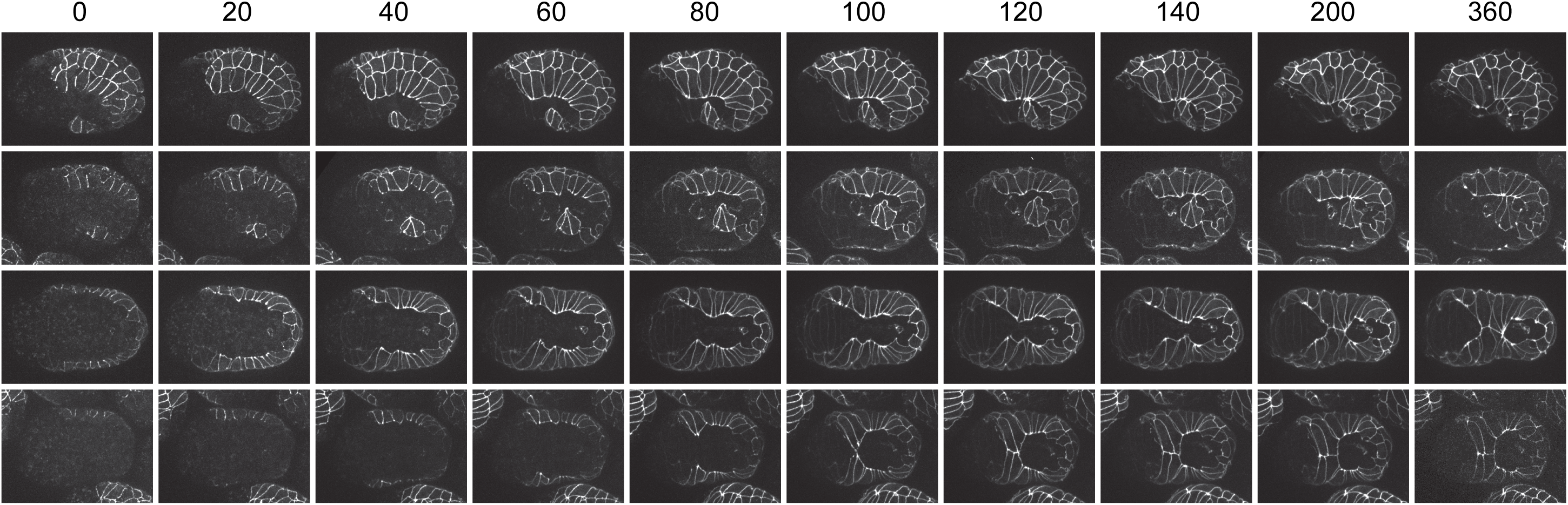
Additional ventral enclosure defects observe in *or1219*ts mutants after late upshifts. Maximum projection images of *or1219*ts mutant embryos expressing DLG-1::GFP to mark epidermal cell membranes after late upshifts. Minutes are listed across the top; time 0 corresponds to the start of imaging at the bean stage (one hour at 26°C after the late upshift time point; see Materials and Methods). Embryos were imaged every 20 minutes for 360 minutes. Each row represents a single embryo over time in the lateral orientation (row 1) and ventral orientation (rows 2-4). The top three rows (1-3) are embryos that exhibited incorrect (non-leading) cell contacts during ventral enclosure. The bottom row (4) shows an embryo that made proper leading cell contact at the ventral midline, but the ventral pocket remained.

## SUPPLEMENTAL TABLES

**Supplemental Table 1.** Strain names for the 191 TS-EL mutant alleles.

| Strain*<br>(original genetic background) | Strain<sup>1</sup><br>(after N2 outcross 1x) | TS allele | Affected<br>gene |
| --- | --- | --- | --- |
| EU847 | EU3058<sup>2</sup> | *or388* | *let-19* |
| EU851 | EU3142<sup>2</sup> | *or392* | *klp-18* |
| CZ4506<sup>1</sup> | EU3136<sup>2</sup> | *or428* | *glp-1* |
| CZ4505<sup>1</sup> | EU3137<sup>2</sup> | *or448* | *glp-1* |
| CZ4115<sup>1</sup> | EU3138<sup>2</sup> | *or488* | *glp-1* |
| CZ4077<sup>1</sup> | EU3065<sup>2</sup> | *or542* | *chaf-1* |
| CZ4385<sup>1</sup> | EU3055<sup>2</sup> | *or566* | *let-19* |
| EU1120 | EU3089<sup>2</sup> | *or614* | *zwl-1* |
| EU1631 | EU2974 | *or789* | *emb-5* |
| EU1632 |  | *or790* |  |
| EU1636 |  | *or794* |  |
| EU1638 | EU3009 | *or796* |  |
| EU1647 | EU3175 | *or805* |  |
| EU1655 | EU3110 | *or813* |  |
| EU1688 |  | *or826* |  |
| EU1676 |  | *or834* |  |
| EU1681 | EU2994 | *or839* | *sart-3* |
| EU1693 |  | *or851* |  |
| EU1695 |  | *or853* |  |
| EU1706 | EU2995 | *or862* | *emb-5* |
| EU1711 | EU3139<sup>2</sup> | *or867* |  |
| EU1713 |  | *or869* |  |
| EU1714 | EU3039 | *or870* | *klp-18* |
| EU1717 | EU3064 | *or873* |  |
| EU1720 | EU2973 | *or876* |  |
| EU1723 |  | *or879* |  |
| EU1727 | EU3111 | *or883* |  |
| EU1737 |  | *or893* |  |
| EU1738 |  | *or894* |  |
| EU1739 |  | *or895* |  |
| EU1740 | EU3040 | *or896* |  |
| EU1742 |  | *or898* |  |
| EU1750 | EU3193 | *or906* |  |
| EU1751 |  | *or907* |  |
| EU1753 |  | *or909* |  |
| EU1758 | EU3001 | *or914* | *gad-1* |
| EU1763 | EU2955 | *or919* | *mom-4* |
| EU1765 | EU3217<sup>2</sup> | *or921* | *nap-1* |
| EU1766 | EU3176 | *or922* |  |
| EU1780 | EU2954 | *or936* | *cdc-25.2* |
| EU1795 |  | *or951* |  |
| EU1798 |  | *or954* |  |
| EU1802 | EU3011 | *or958* |  |
| EU1808 | EU3041 | *or960* |  |
| EU1815 |  | *or967* |  |
| EU1816 |  | *or968* |  |
| EU1822 |  | *or974* |  |
| EU1823 | EU3086<sup>2</sup> | *or975* | *emb-5* |
| EU1829 |  | *or981* |  |
| EU1832 |  | *or984* |  |
| EU1839 | EU3053 | *or991* |  |
| EU1843 | EU3002 | *or995* |  |
| EU1850 | EU3163<sup>2</sup> | *or1002* | *zim-3* |
| EU1851 |  | *or1003* |  |
| EU1852 | EU3042 | *or1004* |  |
| EU1853 | EU3177 | *or1005* | *let-19* |
| EU1854 | EU3194 | *or1006* |  |
| EU1856 |  | *or1008* |  |
| EU1865 |  | *or1016* |  |
| EU1837 |  | *or1024* |  |
| EU1874 |  | *or1025* |  |
| EU1882 |  | *or1033* |  |
| EU1883 |  | *or1034* |  |
| EU1899 | EU2958 | *or1050* |  |
| EU1901 |  | *or1052* |  |
| EU1913 | EU3211 | *or1064* |  |
| EU1920 |  | *or1071* |  |
| EU1931 |  | *or1081* |  |
| EU1933 | EU3012 | *or1083* |  |
| EU1935 | EU2992 | *or1085* | *lrr-1* |
| EU1944 |  | *or1094* |  |
| EU1947 | EU2961 | *or1097* | *emb-5* |
| EU1948 |  | *or1098* |  |
| EU1951 |  | *or1101* |  |
| EU1956 | EU2849 | *or1106* | *zim-3* |
| EU1957 | EU3204 | *or1107* |  |
| EU1958 | EU2963 | *or1108* | *drsh-1* |
| EU1959 | EU3043 | *or1109* |  |
| EU1960 |  | *or1110* |  |
| EU1963 | EU2957 | *or1113* | *let-19* |
| EU1964 |  | *or1114* |  |
| EU1965 |  | *or1115* |  |
| EU1968 | EU2968 | *or1118* | *apx-1* |
| EU1971 | EU3072 | *or1121* |  |
| EU1972 | EU3195 | *or1122* |  |
| EU1976 |  | *or1126* |  |
| EU1982 |  | *or1132* |  |
| EU1987 |  | *or1136* |  |
| EU1996 | EU2993 | *or1140* |  |
| EU2013 | EU3003 | *or1145* |  |
| EU2016 | EU3044 | *or1148* |  |
| EU2021 | EU3212 | *or1153* |  |
| EU2022 | EU2976 | *or1154* | *emb-5* |
| EU2023 |  | *or1155* |  |
| EU2031 |  | *or1163* |  |
| EU2032 |  | *or1164* |  |
| EU2049 | EU2979 | *or1181* |  |
| EU2051 |  | *or1183* |  |
| EU2054 |  | *or1186* |  |
| EU2060 | EU3196 | *or1192* |  |
| EU2061 | EU2841 | *or1193* | *rib-1* |
| EU2063 | EU2972 | *or1195* |  |
| EU2065 |  | *or1197* |  |
| EU2066 | EU3143 | *or1198* |  |
| EU2070 | EU3133 | *or1202* |  |
| EU2080 |  | *or1212* |  |
| EU2082 | EU3112 | *or1214* |  |
| EU2084 | EU3090 | *or1216* |  |
| EU2086 | EU3213 | *or1218* |  |
| EU2087 | EU3148 | *or1219* |  |
| EU2089 |  | *or1221* |  |
| EU2090 | EU3178 | *or1222* |  |
| EU2097 |  | *or1229* |  |
| EU2099 | EU2977 | *or1231* |  |
| EU2105 | EU3013 | *or1237* | *fntb-1* |
| EU2110 |  | *or1242* |  |
| EU2125 |  | *or1257* |  |
| EU2127 | EU3149 | *or1259* |  |
| EU2132 |  | *or1264* |  |
| EU2154 |  | *or1286* |  |
| EU2158 |  | *or1290* |  |
| EU2162 | EU3192 | *or1294* |  |
| EU2164 |  | *or1296* |  |
| EU2167 |  | *or1299* |  |
| EU2168 |  | *or1300* |  |
| EU2169 |  | *or1301* |  |
| EU2170 |  | *or1302* |  |
| EU2177 |  | *or1309* |  |
| EU2180 | EU2843 | *or1312* | *hlh-1* |
| EU2182 |  | *or1314* |  |
| EU2198 | EU3218<sup>2</sup> | *or1330* | *emb-4* |
| EU2206 |  | *or1338* |  |
| EU2210 |  | *or1342* |  |
| EU2222 | EU3113 | *or1354* |  |
| EU2263 |  | *or1395* |  |
| EU2272 | EU3207<sup>2</sup> | *or1404* |  |
| EU2275 |  | *or1407* |  |
| EU2292 | EU3046 | *or1424* |  |
| EU2294 |  | *or1426* |  |
| EU2295 |  | *or1427* |  |
| EU2303 |  | *or1435* |  |
| EU2341 |  | *or1473* |  |
| EU2356 |  | *or1488* |  |
| EU2379 | EU3114 | *or1511* |  |
| EU2400 |  | *or1524* |  |
| EU2402 |  | *or1526* |  |
| EU2415 |  | *or1539* |  |
| EU2416 |  | *or1540* |  |
| EU2478 |  | *or1599* |  |
| EU2481 |  | *or1602* |  |
| EU2531 |  | *or1647* |  |
| EU2552 |  | *or1665* |  |
| EU2554 |  | *or1667* |  |
| EU2555 |  | *or1668* |  |
| EU2559 | EU3063 | *or1672* |  |
| EU2574 | EU2844 | *or1688* | *rib-2* |
| EU2575 |  | *or1689* |  |
| EU2576 |  | *or1690* |  |
| EU2585 |  | *or1699* |  |
| EU2588 | EU3165 | *or1702* |  |
| EU2592 |  | *or1706* |  |
| EU2610 | EU3116 | *or1723* | *emb-9* |
| EU2616 |  | *or1729* |  |
| EU2620 | EU3198 | *or1733* |  |
| EU2624 |  | *or1737* |  |
| EU2625 |  | *or1738* |  |
| EU2626 |  | *or1739* |  |
| EU2627 |  | *or1740* |  |
| EU2629 | EU3015 | *or1742* |  |
| EU2639 | EU3199 | *or1752* |  |
| EU2641 | EU3134 | *or1754* |  |
| EU2642 | EU3078 | *or1755* |  |
| EU2643 |  | *or1756* |  |
| EU2645 | EU3181 | *or1758* |  |
| EU2650 | EU3147 | *or1763* |  |
| EU2652 | EU2839 | *or1765* |  |
| EU2653 | EU3200 | *or1766* |  |
| EU2668 | EU3135 | *or1781* |  |
| EU2669 |  | *or1782* |  |
| EU2671 |  | *or1784* |  |
| EU2672 |  | *or1785* |  |
| EU2674 |  | *or1787* |  |
| EU2678 |  | *or1791* |  |
| EU2684 |  | *or1797* |  |
| EU2687 |  | *or1800* |  |
| EU2733 |  | *or1835* |  |
| EU2787 |  | *or1889* |  |
| EU2792 |  | *or1894* |  |
| EU2793 |  | *or1895* |  |
| EU2811 |  | *or1911* |  |
| EU2832 |  | *or1932* |  |

\* = original genetic background includes: *lin-2(e1309)X*; *ojls1[pie-1::gfp::beta-tbb-2]*; *ruls32[unc-119(+)* *pie-1::GFP::H2B]*; *?ls?[?::gfp::ph]* (may contain *unc-119 (eds)*) III
1 = after N2 outcross 1x, the new strain may contain the following: *ojls1[pie-1::gfp::beta-tbb-2]*; *ruls32[unc-119(+)* *pie-1::GFP::H2B]*; *?ls?[?::gfp::ph]* (may contain *unc-119 (eds)*) III
2 = after N2 outcross 2x, the new strain may contain the following: *ojls1[pie-1::gfp::beta-tbb-2]*; *ruls32[unc-119(+)* *pie-1::GFP::H2B]*; *?ls?[?::gfp::ph]* (may contain *unc-119 (eds)*) III

**Supplemental Table 2.** Strains used for complementation tests (alphabetical by gene).

| Genotype | Strain | Available |
| --- | --- | --- |
| *apx-1(or545ts)* V | EU1013 | BB Lab |
| *cdc-25.2(g52ts)* V | GG52 | CGC |
| *chaf-1(n5453)* I/hT2 [*bli-4(e937)* *let-?(q782)* *qls48*] (I;III) | MT20434 | CGC |
| *dcr-1(ok247)* III/hT2 [*bli-4(e937)* *let-?(q782)* *qls48*] (I;III) | PD8753 | CGC |
| *drsh-1(ok369)* I/ hT2 [*bli-4(e937)* *let-?(q782)* *qls48*] (I;III) | VC1138 | CGC |
| *emb-4(hc60ts)* V | MJ60 | CGC |
| *emb-5(hc61ts)* III | GS1369 | CGC |
| *emb-9(hc70ts)* III | MJ70 | CGC |
| *fntb-1(ok590)* V/nT1 [*qls51*] (IV;V) | VC399 | CGC |
| *gad-1(ct226ts)* *dpy-11(e224)* V | BW1943 | CGC |
| *gad-1(ok573)* V/nT1 (IV;V) | VC327 | CGC |
| *glp-1(q46)* III/ hT2 [*bli-4(e937)* *let-?(q782)* *qls48*] (I;III) | JK4862 | CGC |
| *hlh-1(cc561ts)* II | PD4605 | CGC |
| *klp-18(ok2519)* IV/nT1 [*qls51*] (IV;V) | VC1915 | CGC |
| *let-19(os33)/mln1* [*dpy-10(e128)* *mls14*] II | HS458 | CGC |
| *lrr-1(ok3435)/mln1* [*mls14* *dpy-10(e128)*] II | VC2646 | CGC |
| *dpy-5(e61)* *mom-4(or39)* I/hT1 (I;III); *him-5(e1490)* V | EU424 | BB Lab |
| *mua-3(rh195ts)* III | OT136 | CGC |
| *nap-1(tm1278)* IV/nT1 [*qls51*] (IV;V) | EU1626 | NBRC |
| *pab-1(ok1656)* I/hT2 [*bli-4(e937)* *let-?(q782)* *qls48*] (I;III) | VC1191 | CGC |
| *rib-1(ok556)* IV/nT1 (IV;V) | VC302 | CGC |
| *rib-2(gk318)* III/hT2 [*bli-4(e937)* *let-?(q782)* *qls48*] (I;III) | VC733 | CGC |
| *rpn-1(ok2259)* IV/nT1 [*qls51*] (IV;V) | VC1720 | CGC |
| *sart-3(tm6688)* IV/nT1 (IV;V) | FX16561 | NBRP |
| *tag-307(gk418)* III/mT1 [*dpy-10(e128)*] (II;III) | VC890 | CGC |
| *teg-4(ok883)* I/hT2 [*bli-4(e937)* *let-?(q782)* *qls48*] (I;III) | VC658 | CGC |
| *ttn-1(gk171)* V/nT1 (IV;V) | VC313 | CGC |
| *vab-10(mc44)/unc-75(e950)* *unc-101(m1)* I | ML653 | CGC |
| *zfp-1(ok554)* III | RB774 | CGC |
| *unc-24(e138)* *zim-3(tm2303ts)* IV | CA448 | CGC |
| *zwl-1(ok2378)* I/hT2 [*bli-4(e937)* *let-?(q782)* *qls48*] (I;III) | VC2705 | CGC |

BB Lab = Bruce Bowerman's laboratory
CGC = Caenorhabditis Genetics Center, University of Minnesota, Twin Cities, Minnesota
NBRP = National Bioresource Project, Department of Physiology, Tokyo women's Medical University School of Medicine, Tokyo, Japan

**Supplemental Table 3.**
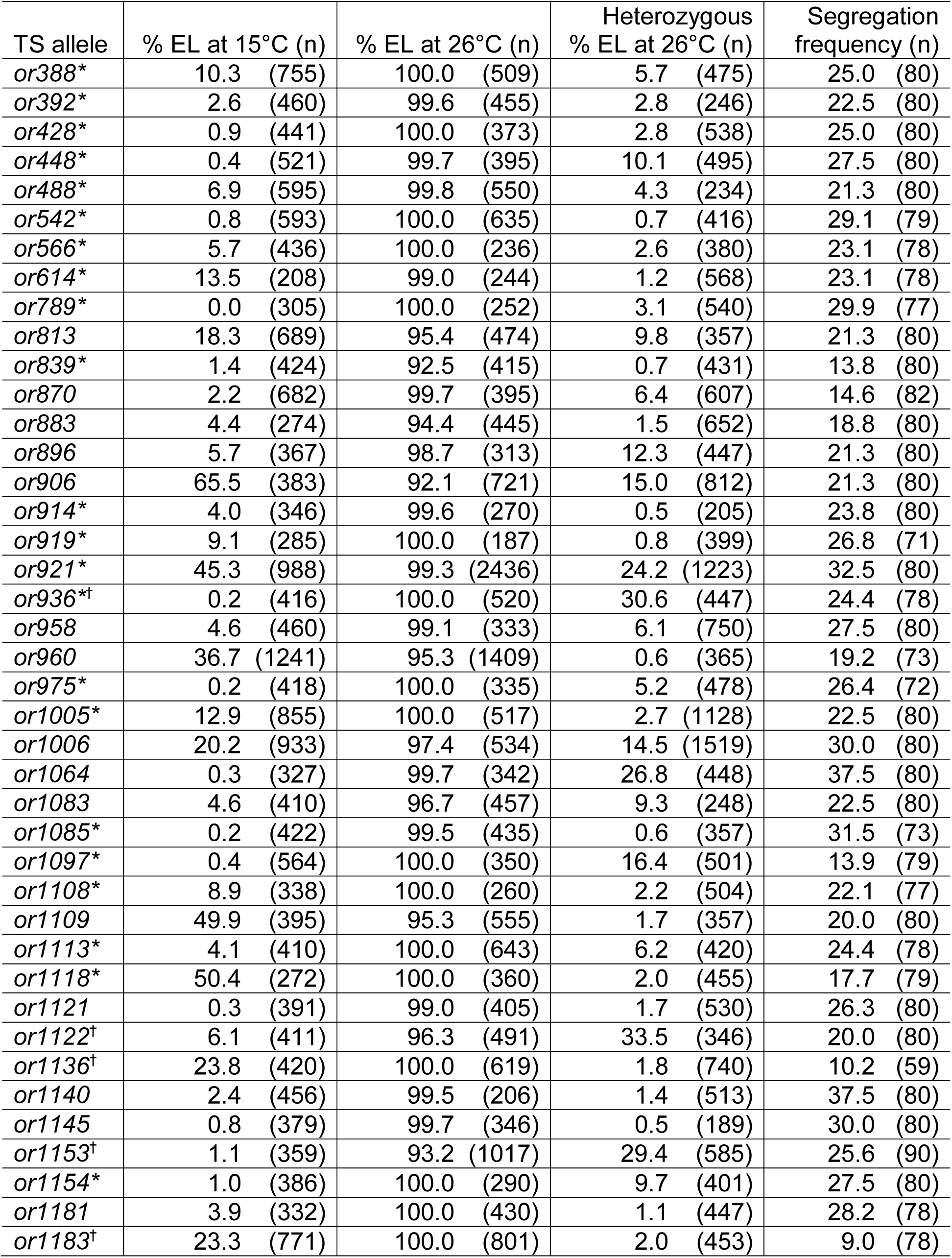
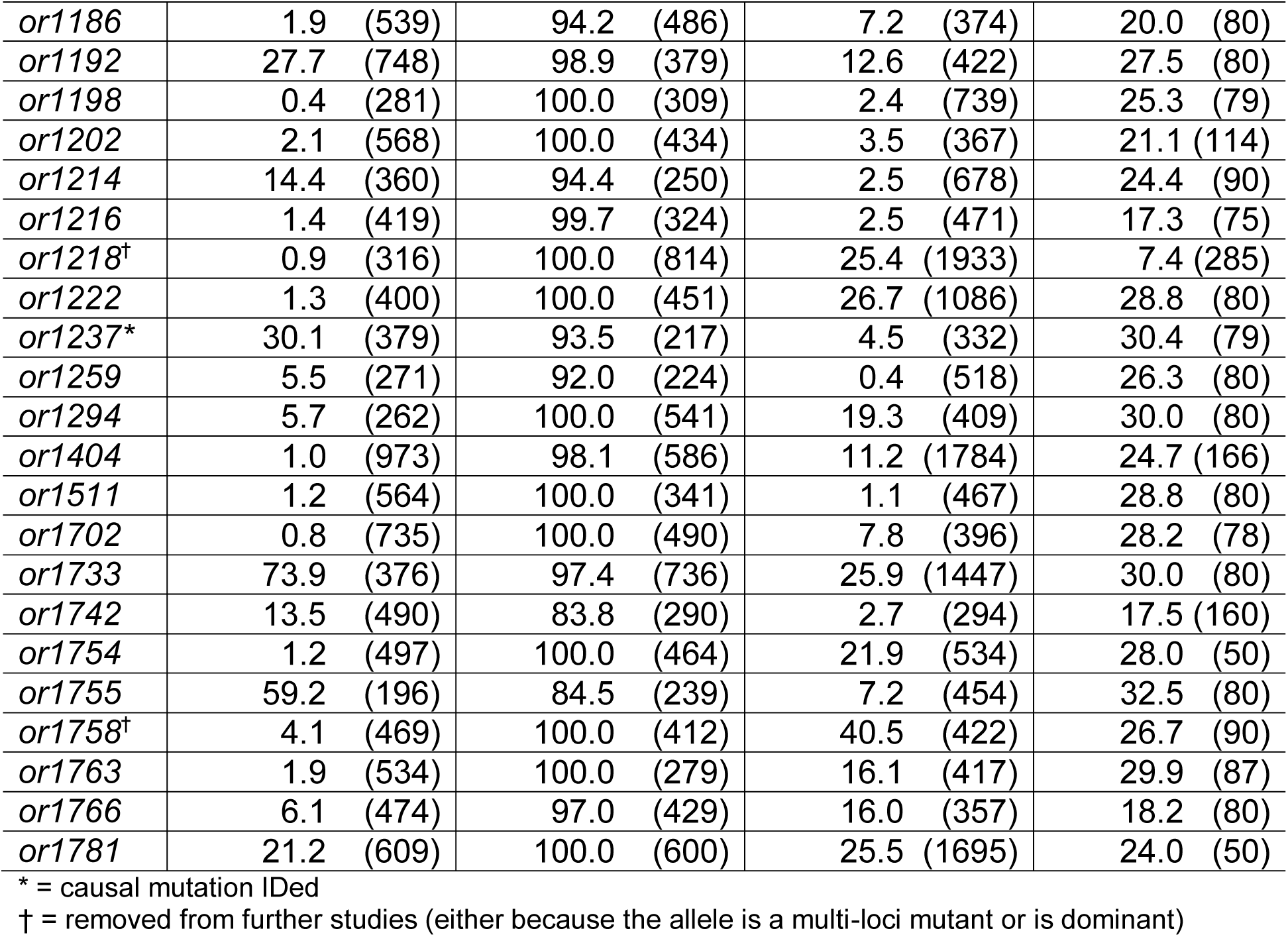
Embryonic lethality and genetic characterization of penetrant L4 upshifted TS-EL mutants.

**Supplemental Table 4.**
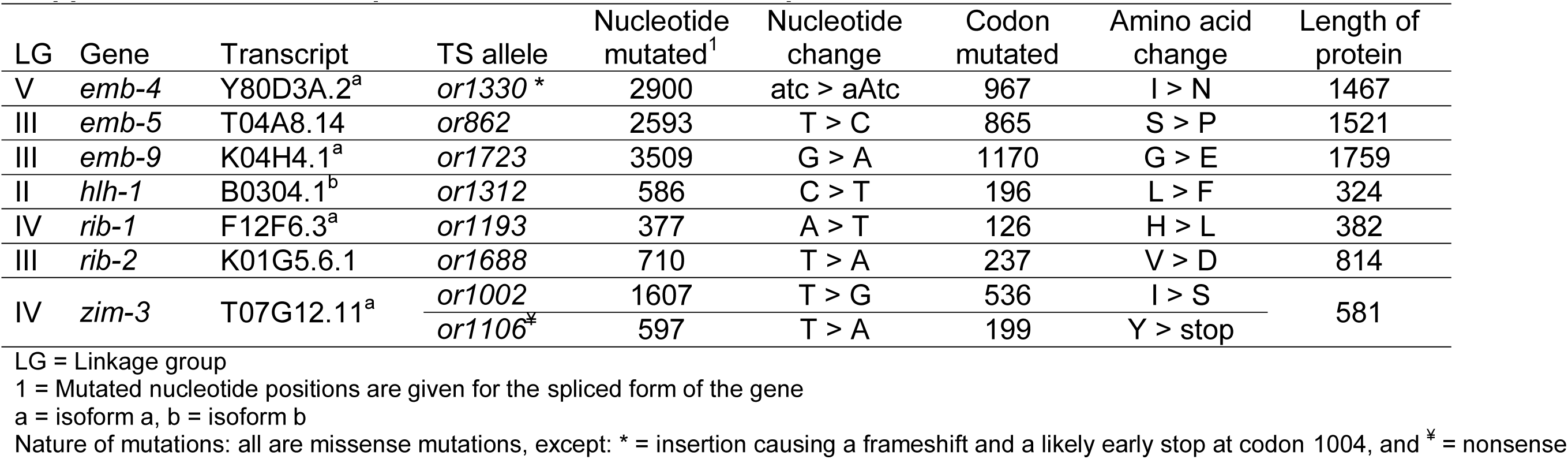
Sequence alterations in the late upshifted causal TS-EL mutants.

**Supplemental Table 5.**
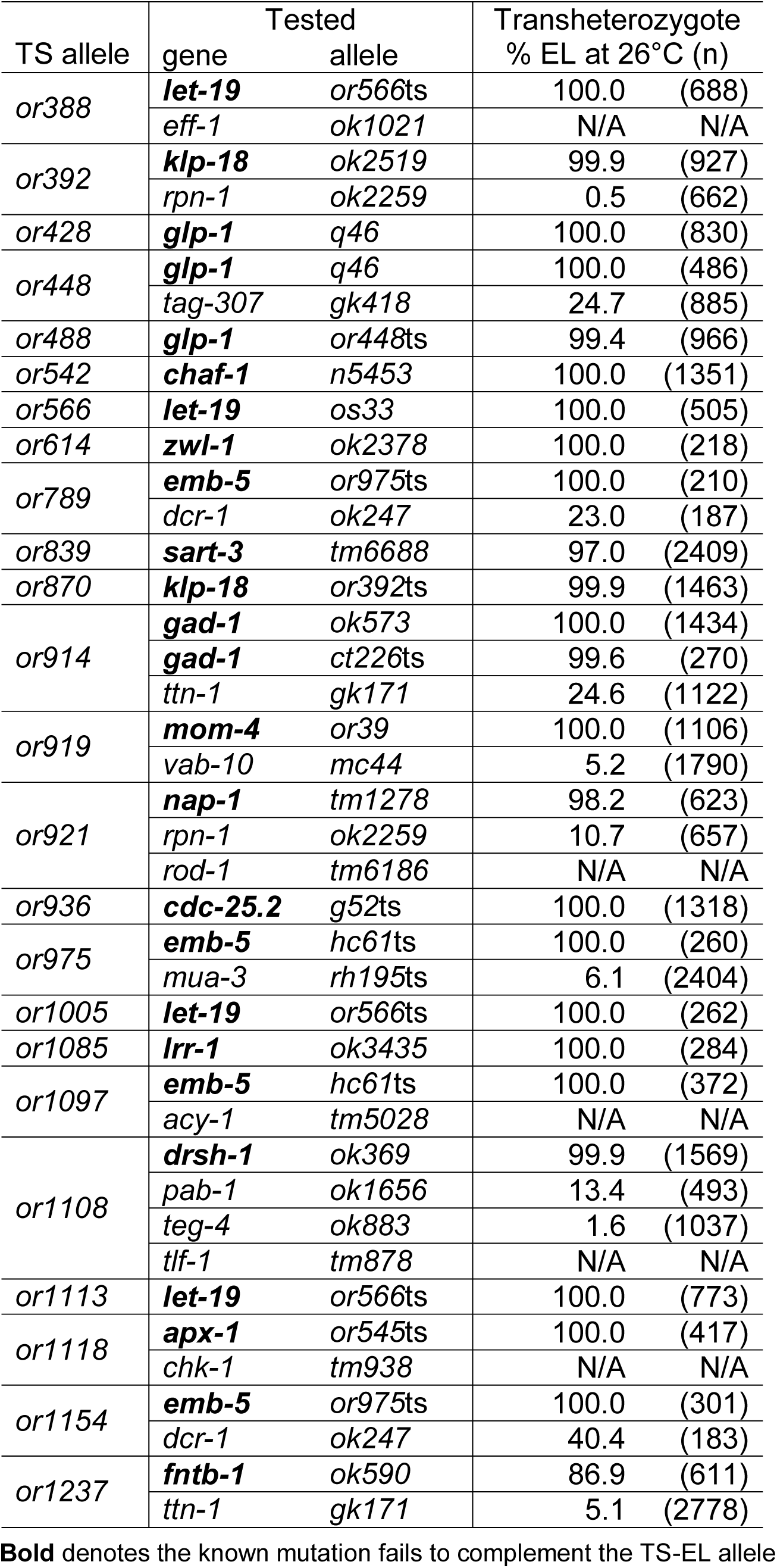
Causal mutation identification for penetrant L4 upshifted TS-EL mutants.

**Supplemental Table 6.**
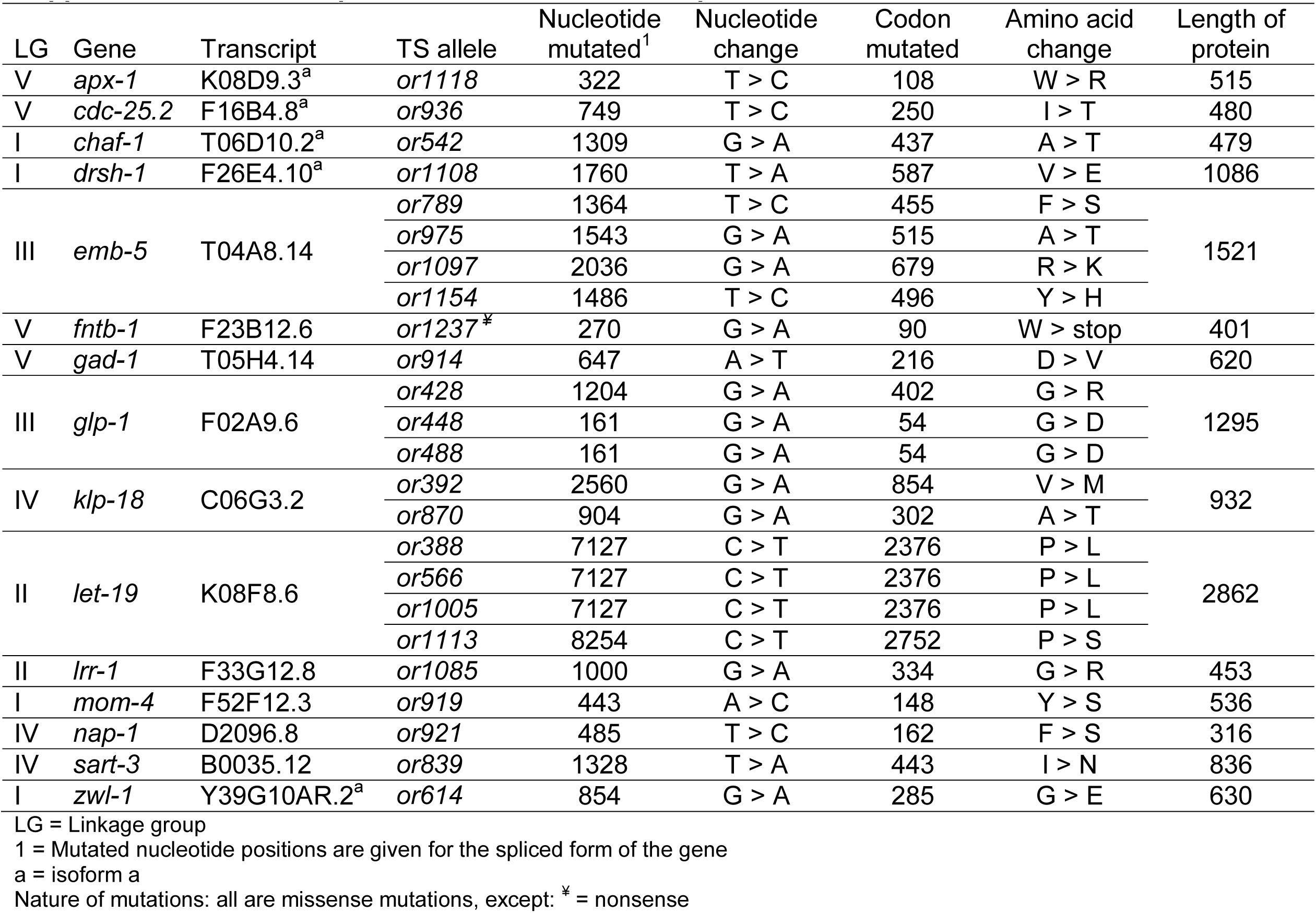
Sequence alterations in the L4 upshifted causal TS-EL mutants.

**Supplemental Table 7.** Gene orthologs and functions for L4 upshifted causal TS-EL mutations.

| Gene | TS allele | Human ortholog | Function |
| --- | --- | --- | --- |
| **Cell Division** (meiosis and/or mitosis) |  |  |  |
| *klp-18* | *or392*<br>*or870* | KLP2 | Kinesin motor protein |
| *zwl-1* | *or614* | ZWILCH | Kinetochore protein |
| **Cell Cycle** |  |  |  |
| *cdc-25.2* | *or936* | CDC25 | Phosphatase |
| *lrr-1* | *or1085* | LRR1 | Substrate recognition subunit of the cullin 2-RING E3 ubiquitin ligase CLR2 |
| **Gene Expression** (signaling/transcription factors) |  |  |  |
| *apx-1* | *or1118* | Delta-like protein1 | Ligand for Notch signaling receptors |
| *emb-5* | *or789*<br>*or975*<br>*or1097*<br>*or1154* | SPT6 | RNA polymerase II transcription elongation factor |
| *gad-1* | *or914* | WDR70 | G protein, protein-protein interactions |
| *glp-1* | *or428*<br>*or448*<br>*or488* | Notch | Transmembrane receptor |
| *let-19* | *or388*<br>*or566*<br>*or1005*<br>*or1113* | TRAP240 | Transcriptional co-activation subunit |
| *mom-4* | *or919* | MAP3K7 | Map kinase |
| **Gene Expression** (nucleosome/chromatin regulation) |  |  |  |
| *chaf-1* | *or542* | CHAF1A | Chromatin assembly factor subunit |
| *nap-1* | *or921* | NAP1L1 | Nucleosome assembly protein |
| **RNA Modification** |  |  |  |
| *drsh-1* | *or1108* | Drosha | RNase III endoribonuclease |
| *sart-3* | *or839* | SART3 | RNA splicing |
| **Protein Modification** |  |  |  |
| *fntb-1* | *or1237* | FNTB | Farnesyltransferase, beta subunit |

